# Minute-scale single-cell transcriptomics reveals a temporal grammar of inflammatory responses

**DOI:** 10.1101/2025.11.15.688181

**Authors:** Kanishk Asthana, Alexander N. Jambor, Wei Wang

## Abstract

Inflammatory stimulation triggers transient pulses, sustained activation, and repression, yet the principles organizing these distinct transcriptional kinetics remain unclear. We developed ChronoSeq, an automated microfluidic platform combining continuous live-cell sampling with molecular time barcoding for pooled, genome-wide single-cell transcriptomics at 10-minute resolution. Across distinct inflammatory contexts, gene expression trajectories repeatedly resolved into a limited set of conserved kinetic modes, despite substantial differences in their timing and gene composition. These modes were associated with distinct functional and regulatory features: promoter architecture was most informative of induction magnitude, whereas RNA turnover features were more closely linked to response persistence and attenuation. Dense temporal profiling further revealed divergent TNF-induced NF-κB responses associated with pre-existing cellular states and resolved delayed inflammatory signaling from LPS-stimulated THP-1 cells to K562 cells. By incorporating temporal ordering into cell–cell communication analysis, we prioritized candidate ligand–receptor interactions whose timing was compatible with downstream activation. Integrating stimulus context with transcriptional dynamics also improved the recovery of known regulatory interactions, particularly for transcription factors whose activity is poorly reflected by their transcript abundance. Together, ChronoSeq reveals an underlying temporal logic of cellular responses, connecting minute-scale transcriptional dynamics to the regulatory architecture and signaling processes that shape them.

## Introduction

Cellular responses to environmental stimuli emerge through dynamic regulatory processes that unfold over minutes to hours.^1–3^ Activation of signaling pathways, transcription factor translocation, transcript turnover, and intercellular communication are all intrinsically temporal phenomena, yet most genome-wide measurements capture them only as static snapshots.^4–13^ As a result, many fundamental questions remain difficult to address: how rapidly do cell states diverge after stimulation, how are signals propagated between cells, and what regulatory principles govern the timing and magnitude of a transcriptional response?

Single-cell RNA sequencing (scRNA-seq) has transformed our ability to characterize cellular heterogeneity, but its application to temporal biology remains limited.^4–13^ Dense time courses are labor-intensive, susceptible to between-timepoint technical variation, and constrained by the physical latency of sample collection, processing and fixation. Consequently, transcriptional responses that occur on rapid timescales are often undersampled, obscuring transient regulatory programs and limiting the reconstruction of accurate cellular trajectories.

Existing experimental platforms provide insights into transcriptional dynamics, but remain limited in their ability to densely sample genome-wide transcriptional responses at minute-scale resolution. Metabolic-labeling assays^1,2^ such as TimeLapse-seq^3^, scSLAM-seq^4^, scNT-seq^5^, and Dyna-vivo-seq^6^ distinguish newly synthesized from pre-existing RNA, enabling the inference of RNA synthesis and degradation rates through modeling. Nascent-transcription assays like GRO-seq^7^, PRO-seq^8^, ChRO-seq^9^, and scGRO-seq^10^ utilize nuclear run-on to capture RNA synthesized from engaged polymerases. Neither approach, however, readily scales to the dense temporal sampling needed to faithfully reconstruct transcriptional responses over time. While Live-seq^11^ enables the serial transcriptomic biopsy of individual living cells, the current implementation is throughput-limited and sub-hour sampling of the same cell has not been demonstrated. In contrast, live-cell imaging^12–14^ can resolve transcription on second-to-sub-minute timescales, but only for a few engineered loci rather than the whole transcriptome.^15^ While highly multiplexed^16^ or robotically batched scRNA-seq could, in principle, support dense time courses, the effective temporal resolution is still likely limited by sampling latency, fixation or quenching time, and asynchronous handling across samples^17,18^. Furthermore, conventional droplet-based workflows require separate sample processing and library preparation across timepoints, making densely sampled, highly multiplexed time courses logistically and financially challenging. Together, these limitations highlight the need for an automated platform capable of genome-wide, minute-scale single-cell profiling under physiological conditions, with broad utility for biological discovery and quantitative modeling of cellular dynamics.

To bridge this gap, we introduce ChronoSeq, an automated microfluidic platform for genome-wide single-cell RNA sequencing with sampling intervals as brief as ten minutes. ChronoSeq integrates continuous live-cell sampling with molecular time-barcoding in a unified workflow, enabling densely timed transcriptional profiling while preserving cellular health. We further developed ChronoPack, a computational package for reconstructing cellular trajectories, modeling transcriptional kinetics, inferring gene regulation, and analyzing intercellular communication from densely sampled single-cell time courses.

Using inflammatory signaling as a model system, we show that minute-scale transcriptomic profiling reveals response heterogeneity and intercellular signaling dynamics that are otherwise inaccessible to conventional sampling. In TNF-α-stimulated K562 cells, ChronoSeq resolved two distinct response trajectories, with a pre-existing erythroid-like subpopulation exhibiting attenuated NF-κB responsiveness. This finding links baseline transcriptional heterogeneity to variation in the magnitude and kinetics of inflammatory activation. In an LPS-stimulated THP-1–K562 coculture, we further resolved the temporal progression of a delayed paracrine response, implicating TNF as a candidate mediator based on the timing of its induction relative to downstream K562 activation.

Across these cellular and signaling contexts, we identified a conserved repertoire of transient, sustained, and repressed transcriptional response modes, even as their gene composition and timing differed. These modes were associated with distinct cis-regulatory and transcript-level features: promoter architecture was most predictive of induction magnitude, whereas RNA stability was more closely associated with response attenuation and persistence. Finally, we demonstrate the value of high-density temporal data for inferring gene regulation in rapid signaling contexts. By applying ChronoPack’s temporally-informed inference, we outperformed established co-expression-based methods and achieved competitive performance relative to perturbation-derived rankings in recovering known regulatory targets of key stimulus-responsive transcription factors.

Together, ChronoSeq and ChronoPack establish an integrated framework for measuring and modeling cellular dynamics at minute-scale resolution. By moving from static transcriptomic snapshots to densely sampled single-cell time courses, this approach lays the foundation for quantitative and predictive models of cellular behavior.

## Results

### An automated platform for minute-scale temporal scRNA-seq

ChronoSeq addresses the sampling latency, fixation artifacts, and between-timepoint batch effects that collectively limit conventional scRNA-seq from resolving minute-scale transcriptional dynamics. Cells are maintained continuously under physiological conditions (37°C, 5% CO₂) throughout the experiment, ensuring that captured profiles reflect authentic biological responses. At programmed intervals as short as seven to ten minutes, the device automatically diverts a defined cell population directly into droplet co-encapsulation with lysis buffer (**Fig. 1a**, **Fig. S2, Supplementary Movie 1**). Immediate lysis upon droplet formation avoids fixation delays and preserves the transcriptional state at the sampling time. Each sampled population is co-encapsulated with a uniquely barcoded Time-Tag bead, allowing all timepoints to be pooled into a single library and processed under identical conditions to dramatically reduce labor while minimizing batch effects and manual variability **(Fig. S1)**.

**Figure 1.**
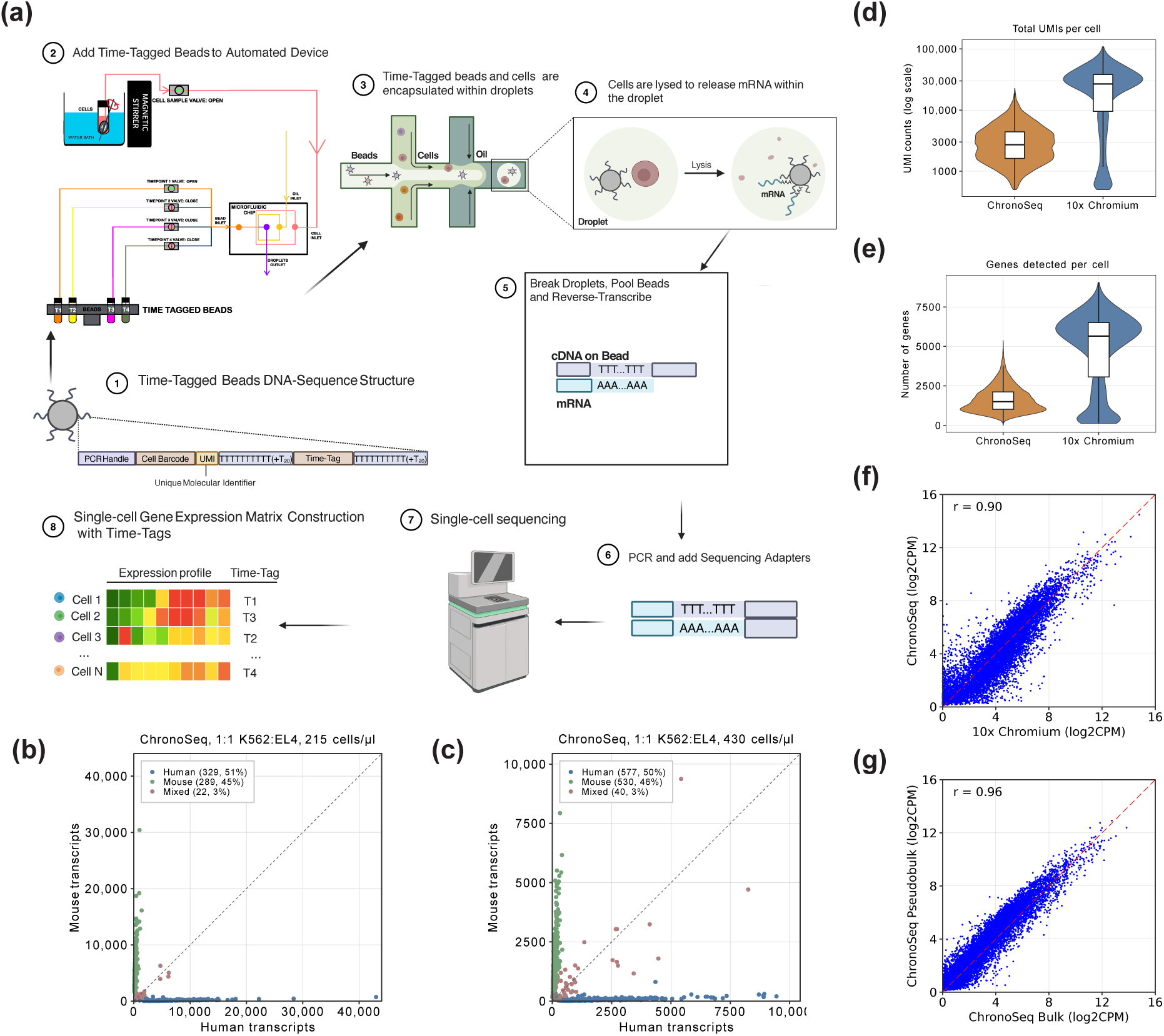
**(a)** *ChronoSeq Library Preparation Workflow*: Time-Tagged beads are added to reservoirs inside the ChronoSeq device (Step 1-2). Our current system supports 12 unique Time-Tags, each Time-Tag is added to its own dedicated reservoir. Cells suspended in a culture media are loaded into another reservoir maintained at 37°C and 5% CO2. The ChronoSeq device then automatically co-injects Time-Tags 1-12 with samples 1-12 from the cell reservoir into a microfluidic chip at programmed time intervals of ≥7 minutes (Supplementary Movie 1 and Figure S2), generating droplets in which cells from each sampling time point are paired with beads carrying the corresponding Time - Tag (Step 3). Cells are immediately lysed upon droplet formation and kept on ice to preserve the transcriptome until all 12 samples are collected (Step 4). For samples with high RNase activity, RNase inhibitor is added to each sample to help further maintain RNA integrity. Next, following the same process as Drop-seq, droplets are broken, pooled, and reverse transcribed together in unified library preparation workflow for all samples. The cDNA on the beads is amplified and sequencing adapters are added (Step 5-6). The library is sequenced and processed with our ChronoSeq-Tools pipeline to create Digital Gene Express (DGE) matrices with an assigned Time-Tag for each cell (Step 7-8). (**b,c)** Species-mixing barnyard plots for 50:50 K562 (Human) and EL4 (Mouse) cells at two commonly used ChronoSeq working concentrations (215 cells/ul and 430 cells/ul). (**d-g**) We generated K562 negative control datasets using both single-cell and bulk sampling with the ChronoSeq device. For both datasets, K562 cells were maintained inside the ChronoSeq device and sampled at 10-minute intervals. For the bulk dataset, cells sampled by the device were manually mixed with Time-Tags 1-12. **(d-e)** We downloaded both replicates of a published K562 dataset^19^ generated using the commercially available 10X Genomics Chromium 3’ single-cell RNA-seq technology for comparison: **(d)** shows a violin plot comparing median transcripts captured, **(e)** shows a violin plot comparing genes detected for the 10X Genomics dataset and ChronoSeq single-cell K562 negative control dataset. **(f-g)** We prepared pseudobulks across all Time-Tags for both bulk and single-cell ChronoSeq K562 negative control datasets. We also prepared a combined pseudobulk for both 10X Genomics K562 replicates. **(f)** shows a correlation scatterplot between the 10X Genomics and ChronoSeq single-cell K562 pseudobulks. ChronoSeq shows a high concordance with 10X Genomics with a correlation of r=0.90. **(g)** Comparison of ChronoSeq single-cell and bulk shows even higher concordance with a correlation of r=0.96.

To implement this multiplexed workflow, we manufactured (**Fig. S3**) and validated a custom 12-plex Time-Tag barcode system using mixed human–mouse samples. Each tag cleanly partitioned species-specific transcripts, with minimal barcode cross-talk (<=1.6% mixed beads per tag; **Fig. S4**). Automated sampling within the ChronoSeq device showed similarly precise tagging and assigned cells to their intended timepoint barcode (**Fig. S5**). ChronoSeq’s single-cell doublet rate matched commercial standards, at below 4% across operational cell concentrations (**Fig. 1b,c**). We captured an average of 636 cells per Time-Tag. Together, these results demonstrated that ChronoSeq achieves fully automated, high-integrity sample tagging and single-cell encapsulation.

To benchmark transcriptional accuracy independently of temporal resolution, we profiled unstimulated K562 cells and compared ChronoSeq with established references.^19^ Despite the number of detected genes and transcripts per cell being lower than 10X Chromium (**Fig. 1d,e**), ChronoSeq exhibited quantitative fidelity: pseudobulk profiles from ChronoSeq single cells correlated nearly perfectly with ChronoSeq bulk RNA-seq (r = 0.96) and strongly with 10x Chromium pseudobulks (r = 0.90) (**Fig. 1f,g**). These results confirmed that ChronoSeq accurately preserves transcriptional signatures, with minimal bias introduced by automated sampling.

### ChronoPack overcomes the computational challenges of high-density temporal scRNA-seq

ChronoSeq dense temporal sampling introduces analytical challenges that differ fundamentally from those encountered in conventional scRNA-seq. Many trajectory inference approaches rely primarily on transcriptomic similarity to order cells along developmental or response continua, an assumption that is most informative when cellular progression is accompanied by appreciable changes in expression state. At 10-minute resolution, however, adjacent snapshots may be only weakly separated in expression space, making static transcriptional distance an imperfect proxy for temporal progression. Under these conditions, trajectory reconstruction depends on resolving subtle local transitions across consecutive timepoints rather than large global state changes, complicating the detection of transient intermediate states and early response divergence. In parallel, dynamic gene identification becomes statistically challenging because meaningful temporal programs may appear as smooth, transient, or distributed deviations across adjacent measurements rather than large endpoint shifts. Together, these features motivate analytical approaches that explicitly leverage dense temporal continuity rather than relying on static transcriptional similarity alone.

To address these challenges, we developed ChronoPack, a computational framework designed specifically for the analytical demands of densely sampled temporal scRNA-seq (**Fig. 2a**). ChronoPack reconstructs trajectories by modeling cellular dynamics as a sequence of locally constrained optimal transport (OT) problems using WaddingtonOT^20^. This formulation is particularly well matched to dense temporal data: because short inter-timepoint intervals justify the assumption that cells transition minimally between adjacent measurements, the OT problem becomes better conditioned and cellular progression can be reconstructed through local transitions rather than relying on global trajectory assumptions or continuous-time parameterization^21–23^, both of which become unreliable when few distant timepoints are available. From the resulting transition kernel, ChronoPack propagates cells forward through the full time course to estimate probability distributions over terminal states. Trajectory structure is then derived from similarity in these future-state probability profiles rather than from traditional expression-based clustering (see **Methods**).

**Figure 2.**
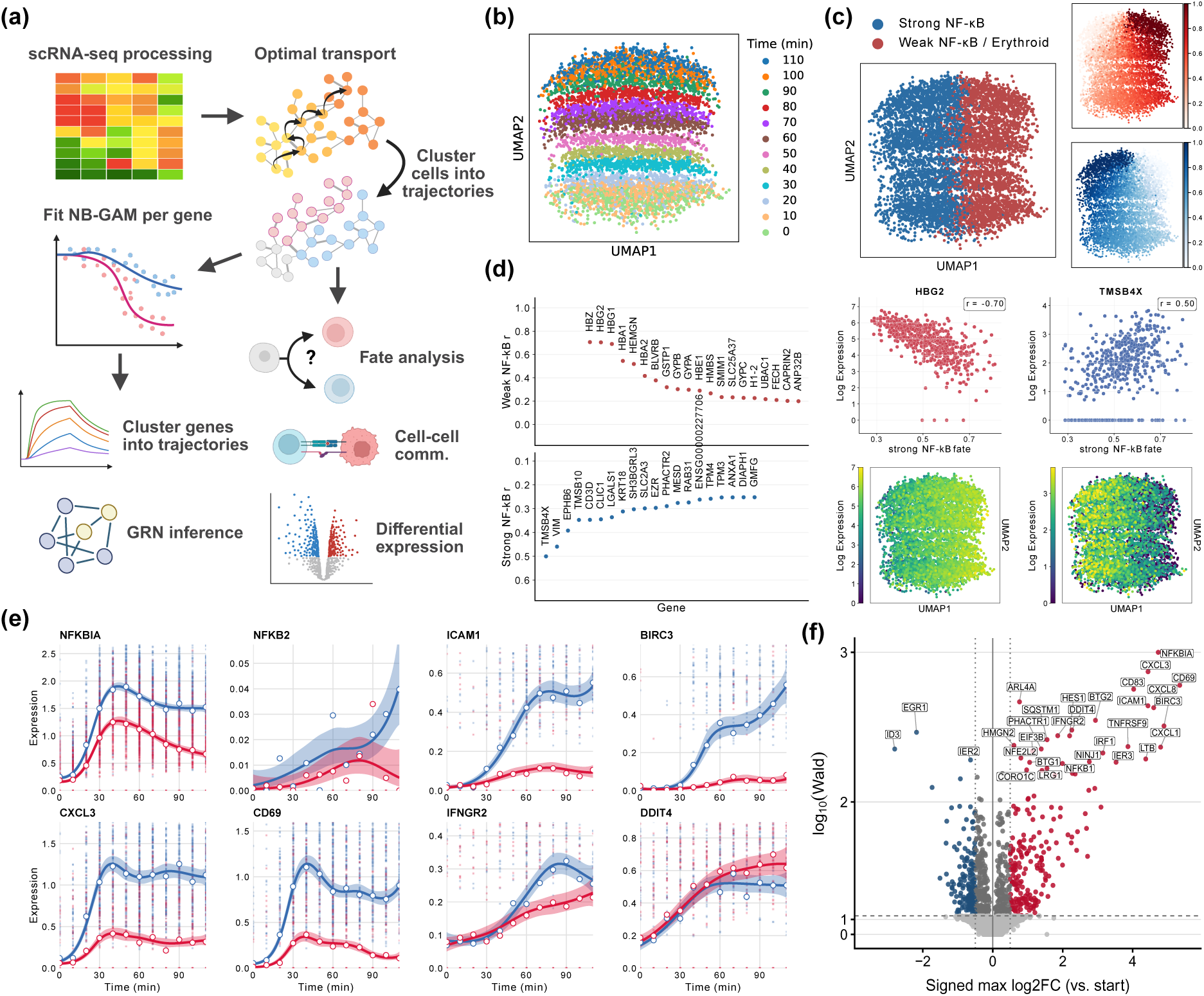
ChronoPack resolves cell-state-dependent NF-κB dynamics in a high-resolution scRNA-seq time course. **(a)** Overview of the ChronoPack workflow. **(b)** UMAP of the K562 scRNA-seq time series, constructed from a WaddingtonOT-derived transition kernel and colored by sampling time. **(c)** Visualization of the strong (non-erythroid) and weak (erythroid-like) NF-κB response trajectory clusters and their corresponding inferred cell fate probabilities. **(d)** Genes associated with divergent NF-κB response outcomes. *(left)* Top genes whose early expression was associated with either fate; *(right)* Representative genes showing positive (TMSB4X) or negative (HBG2) associations with NF-κB response strength. **(e)** Fitted expression trajectories for representative genes, showing attenuated induction of canonical NF-κB targets in erythroid-like K562 cells alongside gene-specific differences in response shape. **(f)** Dynamic gene discovery across the time course. Genes are plotted by signed maximum log2 fold change relative to the initial time point and temporal association strength from tradeSeq; significant induced and repressed genes are highlighted.

ChronoPack integrates this trajectory structure into a unified set of downstream analyses. Temporally-informed embeddings are generated by applying UMAP to a symmetrized graph constructed from the WaddingtonOT transition kernel, producing visual layouts that reflect inferred temporal connectivity rather than expression similarity alone. Genome-wide kinetic modeling within each trajectory uses negative-binomial generalized additive models (NB-GAMs)^24^ to detect smooth temporal expression dynamics while accounting for count noise and overdispersion, enabling detection of gradual, transient, or rapidly decaying responses missed by pairwise endpoint comparisons. These cellular trajectories and gene kinetic profiles enable us to classify transcriptional responses, compare regulatory programs across contexts, and infer cell–cell communication and gene regulatory networks. Together, ChronoPack converts dense temporal scRNA-seq measurements into an interpretable model of dynamic cellular behavior, with each analytical layer building on the inferred temporal structure.

### Pre-existing cell-state heterogeneity is associated with divergent NF-κB responses

To demonstrate the biological utility of ChronoSeq and ChronoPack, we profiled K562 cells responding to TNF-α at 10-minute intervals over two hours. TNF-α stimulation rapidly activates canonical NF-κB signaling through TNFR1/2-mediated nuclear translocation of RELA/p65–NFKB1/p50. Temporal pseudobulk profiles showed strong concordance with matched qPCR measurements for representative NF-κB target genes (**Fig. S6**), confirming quantitative fidelity of the ChronoSeq measurements.

Trajectory reconstruction with ChronoPack resolved two K562 subpopulations with distinct TNF-induced inflammatory response trajectories (**Fig. 2b,c,e**). The larger population mounted a strong inflammatory response, marked by induction of canonical NF-κB targets including *NFKBIA*, *NFKB2*, *ICAM1*, *BIRC3*, and *CXCL3*. By contrast, a smaller erythroid-like subpopulation retained expression of lineage-associated genes such as *HBE1*, *HBA2*, and *GYPC*, but exhibited a substantially attenuated NF-κB response. This attenuation extended across distinct response kinetics: rapidly induced genes such as NFKBIA and CD69 showed weaker and less persistent induction, whereas more gradually accumulating targets such as ICAM1 and BIRC3 remained weakly induced throughout the time course (**Fig. 2e**). Notably, DDIT4 showed greater induction in the erythroid-like subpopulation, indicating that reduced responsiveness was not shared across all induced genes. This population also showed lower expression of *TNFRSF1A*, suggesting that erythroid-like state heterogeneity may preconfigure TNF responsiveness in part through reduced receptor expression. To assess whether these response differences were associated with initial state, we correlated gene expression at the first timepoint with terminal fate probabilities inferred from the transition kernel. This analysis associated higher expression of erythroid genes, including HBG2 (Spearman ρ = −0.71), with weaker TNF-α responsiveness, whereas cytoskeletal-associated genes such as TMSB4X (ρ = 0.50), VIM, and TMSB10 were associated with stronger responses (**Fig. 2d**). Together, these findings suggest that pre-existing state heterogeneity shapes TNF-induced NF-κB responsiveness. Our results extend prior work on hemin-induced erythroid differentiation^25^ by showing that naturally occurring erythroid-biased K562 states are linked to attenuated NF-κB dynamics.

### Genome-wide classification resolves distinct dynamic response modes

Genome-wide ChronoSeq profiling enabled unbiased identification of transcriptional response dynamics beyond static differential expression. Using ChronoPack’s NB-GAM framework, we identified 2,022 genes exhibiting nominally significant temporal dynamics (p < 0.05) among 20,036 expressed genes, including a high-confidence subset of 434 genes (q < 0.05, |log2FC| > 0.5), representing 2.2% of the detected transcriptome. Despite similar proportions of induced and repressed genes (47% versus 53%), induction was associated with substantially larger changes in expression magnitude (**Fig. 2f**). To characterize the response shapes, fitted gene expression trajectories were normalized and clustered into six broad dynamic modes: impulse-up (two subtypes), trend-up, trend-down (two subtypes), and transient-down. (**Fig. 3a**). The impulse-up subtypes differed in peak timing and subsequent decay, whereas the trend-down subtypes were distinguished primarily by trajectory curvature. Each broad dynamic mode comprised several higher-resolution clusters that captured additional variation in trajectory shape. Individual gene trajectories were further characterized using descriptive features, including onset and peak time, response persistence, area-under-the-curve (AUC), and peak log_2_ fold change, providing a quantitative description of each gene’s kinetics.

**Figure 3.**
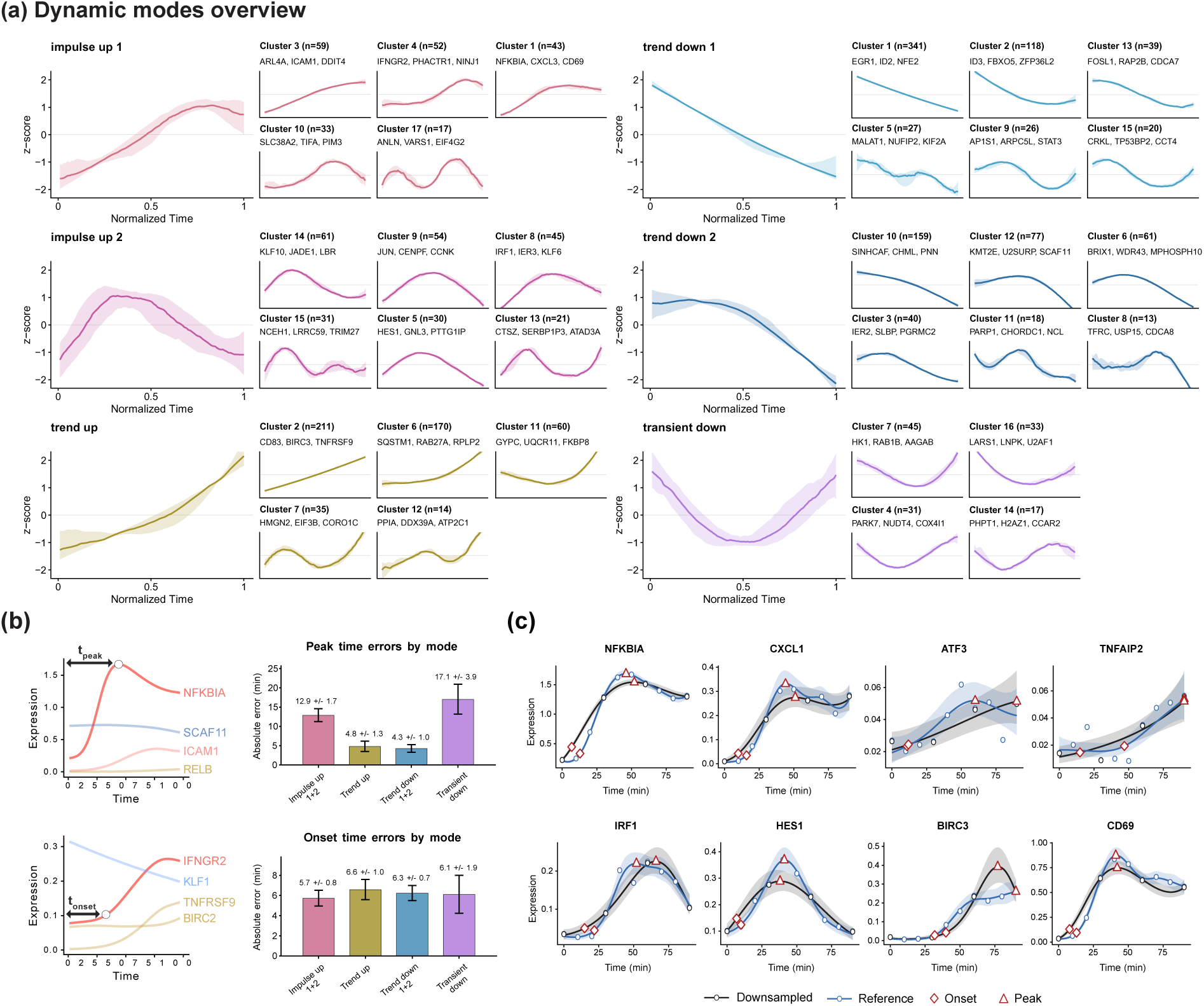
ChronoSeq resolves dynamic response modes and reveals the effects of temporal downsampling. **(a)** Major dynamic response modes identified by hierarchical clustering of fitted gene expression trajectories, together with their constituent clusters and representative genes. **(b)** (*left*) Representative genes illustrating how peak and onset times are calculated. (*right*) Mean absolute differences in estimated peak and onset times between the original 10-minute time course and its synthetically downsampled 30-minute counterpart, stratified by dynamic mode. **(c)** Comparison of gene expression trajectories fitted to the original 10-minute dataset (blue; reference) and the 30-minute downsampled dataset (black). Diamonds and triangles indicate estimated onset and peak times, respectively. Coarser sampling preserves broad expression patterns but can alter estimated timing and obscure finer features of transient responses.

The impulse-up and trend-up modes captured distinct forms of TNF-α-induced transcription. Impulse-up genes showed rapid induction followed by varying degrees of attenuation, whereas trend-up genes increased steadily over time. The impulse-up mode consisted of two kinetic subtypes: a broader, more sustained response, including genes such as *NFKBIA*, *CXCL3*, *CD69*, *ICAM1*, *CXCL8* and *IFNGR2*, and a sharper, more transient response marked by *JUN*, *IRF1*, *HES1*, *KLF6* and *JUNB* (**Fig. S11**). Many of these genes are canonical NF-κB targets and immediate-early inflammatory regulators, with *NFKBIA* serving as a particularly clear example – acting both as a rapidly-induced NF-κB target and as a core negative-feedback regulator that re-sequesters nuclear NF-κB complexes to the cytoplasm. By contrast, trend-up genes, including *RELB*, *TNFRSF9*, *SQSTM1*, *LTB* and *IL27RA*, were induced on a similar timescale but continued to accumulate throughout the measured interval rather than reaching an early peak and attenuating or plateauing. Together, these trajectories show that TNF-responsive genes follow distinct induction kinetics, ranging from rapid pulses to progressively accumulating responses.

Repressed genes likewise resolved into distinct kinetic programs. The trend-down and transient-down modes captured genes whose expression decreased following TNF-α stimulation, with the latter group showing partial recovery within two hours. Trend-down genes separated into a smooth, monotonic subtype (e.g. *KLF1*, *NFE2*, *FOSL1*, *ZFP36L2* and *GATA1*), and a second subtype marked by greater curvature and delayed repression (e.g. *IER2*, *H2AC17*, *MEF2C*, *KDM5B*). Notably, *KLF1*, *NFE2*, and *GATA1* were all repressed, pointing to the attenuation of the K562 erythroid program during inflammatory activation. More broadly, because NF-κB and AP-1 lack well-established direct repressive functions at many of these loci, these coordinated decreases are unlikely to reflect widespread direct repression; instead, they may arise from ongoing remodeling of the pre-existing cell state, redistribution of regulatory cofactors, or both.

Consistent with this possibility, acute NF-κB activation has been shown to redistribute cofactors such as BRD4 and MED1 toward inflammatory enhancers at the expense of lineage-restricted regulatory elements^26^. ChronoSeq thus resolves transcriptional dynamics consistent with early destabilization of the erythroid regulatory program occurring simultaneously with activation of inflammatory signaling, revealing the temporal relationship between these transcriptional changes.

### Dense temporal sampling improves the reconstruction of rapid transcriptional responses

To assess the impact of sampling resolution on kinetic reconstruction, we synthetically downsampled the K562 time course from 10-minute to 30-minute intervals and repeated trajectory inference and dynamic gene modeling (**Fig. 3b, c**). Reduced sampling density preserved the broad organization of the TNF response but substantially degraded kinetic precision: peak and onset time estimates shifted by an average of 7.9 and 6.1 minutes for 238 high-confidence dynamic genes, respectively, and the transient induction profiles of several early NF-κB targets were reconstructed as broader, shallower trajectories. Notably, subtle rebound-like features in some NF-κB-responsive trajectories were diminished or lost under coarser sampling, making these responses appear more monotonic. Such features are consistent with the complex transcriptional dynamics associated with canonical NF-κB negative feedback, in which newly synthesized IκBα re-sequesters nuclear RELA and can contribute to oscillatory signaling. While these transcript-level patterns do not directly establish NF-κB oscillations, their loss illustrates how sparse sampling can obscure features potentially associated with feedback regulation. These distortions were most pronounced among impulse-class genes, whose sharp transient behavior is particularly sensitive to temporal sampling density. Thus, this comparison establishes minute-scale resolution as a functional requirement, rather than merely a technical refinement, for accurately reconstructing the timing, ordering, and kinetic architecture of fast-acting transcriptional programs. By preserving the temporal structure of transcriptional responses, dense temporal profiling provides a strong foundation for mechanistic modeling and extends beyond differential expression to a more faithful reconstruction of inflammatory gene expression dynamics.

### High-density temporal profiling resolves the timing of intercellular signaling

To test whether ChronoSeq could resolve dynamic cell-cell communication, we modeled innate immune secondary signaling in a co-culture of LPS-responsive THP-1 macrophages and K562 cells, which showed no appreciable direct LPS response in a separate control setting (**Fig. S7**). Conventional single-cell analyses infer communication primarily from ligand-receptor co-expression, but such approaches cannot determine the temporal ordering or latency of signaling events. In contrast, ChronoSeq’s dense 15-minute sampling directly captured the sequential propagation of an inflammatory signal across cell types. Following LPS stimulation, THP-1 cells rapidly initiated the canonical inflammatory program, while K562 cells displayed a delayed, secondary response consistent with signaling from THP-1-derived factors (**Fig. 4a, b).** Comparison of transcriptional response amplitudes between THP-1 and K562 (**Fig. 4c**) revealed an LPS/TLR4-biased module (TNFAIP2, CCL3, and CCL4, among other genes) that was distinct from the K562 secondary response (CD69, IRF1, LTB). A subset of chemokines (including CXCL3 and CXCL8) were preferentially but not exclusively expressed in THP-1 cells. These differences underscore the functional distinction between the LPS-induced and paracrine inflammatory responses in THP-1 and K562 cells, respectively.

**Figure 4.**
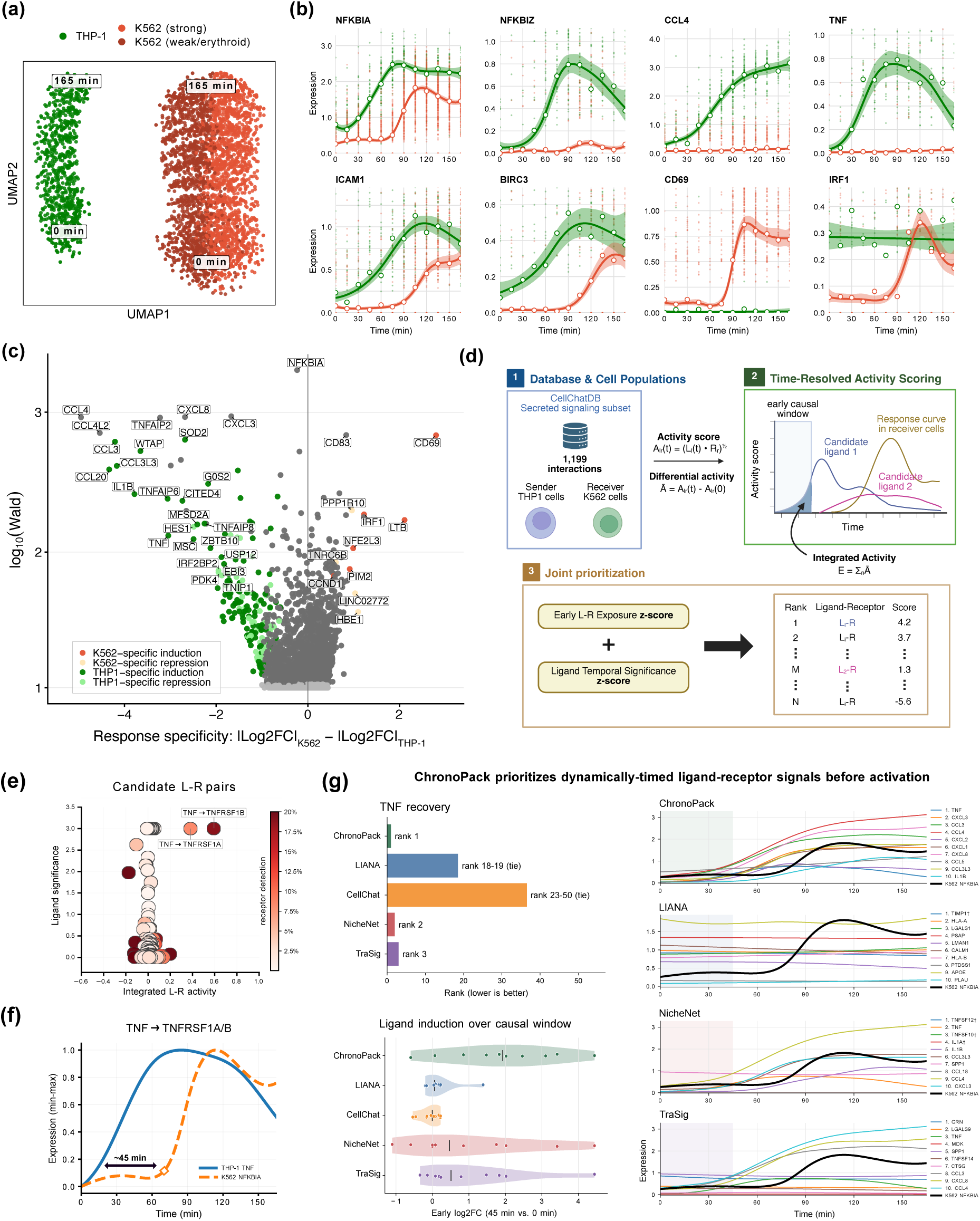
ChronoSeq resolves paracrine TNF–NF-κB signaling and prioritizes candidate ligand–receptor interactions mediating cell-cell communication. LPS was added 5 minutes after the first sample, followed by 15-min sampling across the remaining 11 time points. At this dose, THP-1 cells respond directly through TLR4, whereas K562 cells lack a direct LPS response (**Fig. S7**) and report paracrine inflammatory signaling instead. **(a)** UMAP of THP-1 and K562 cells colored by cell type and K562 NF-κB response trajectory. **(b)** Gene expression trajectories for shared NF-κB response genes alongside stimulus-specific genes. THP-1 shows direct LPS-induced inflammatory activation, whereas K562 cells exhibit delayed NF-κB activation. **(c)** Response specificity of significantly dynamic genes, quantified as the difference in maximum log2 fold change between K562 and THP-1. Highlighted genes mark cell type–specific induction or repression. **(d)** Overview of ChronoPack’s ligand–receptor (L-R) prioritization for THP-1-to-K562 signaling. Candidate interactions from CellChatDB were evaluated using ligand expression dynamics in THP-1 and receptor expression in K562. Time-dependent L-R activity was integrated over a putative causal window preceding K562 activation and combined with ligand temporal significance to score candidate interactions. **(e)** ChronoPack’s L-R prioritization identifies TNF–TNFRSF1A and TNF–TNFRSF1B as top candidates. Axes indicate integrated L–R activity and ligand temporal significance, while color denotes receptor detection fraction. **(f)** Overlay of TNF expression in THP-1 (blue) and NFKBIA expression in K562 (orange), indicating an approximately 45-minute lag between THP-1 TNF and K562 NFKBIA transcript induction, consistent with the proposed paracrine signaling pathway. Expression values are min-max normalized over time. (**g**) Benchmarking of ChronoPack C2C against other cell-cell communication inference tools, including coexpression-based (LIANA, CellChat), pseudotime-dependent (TraSig), and ligand-target-based (NicheNet) methods. (*top left*) Recovery of TNF as the expected operational ligand. (*bottom left*) Induction of predicted ligands within the putative causal window preceding K562 activation, quantified as log2-fold change from 0 to 45 min. (*right*) Expression curves of the top-ranked ligands by each method.

To identify candidate mediators of the delayed K562 response, we quantified time-resolved ligand–receptor (L–R) activity scores between THP-1-expressed ligands and K562-expressed receptors using CellChatDB^28^ annotations, and integrated these scores over a putative causal window preceding K562 activation (**Fig. 4d**). We defined the causal window to occur at least 30 minutes before K562 response onset, accounting for the delay imposed by ligand production, secretion, receptor engagement and downstream signaling. Among the 230 L-R pairs screened, TNF-TNFRSF1A and TNF-TNFRSF1B received the highest combined scores, reflecting early TNF induction together with detectable expression of its cognate receptors in K562 (**Fig. 4e**). These results prioritized TNF as a candidate mediator of the secondary inflammatory response; consistent with this, *TNF* transcripts rose in THP-1 cells immediately following LPS stimulation and preceded induction of NF-κB targets in K562 by approximately 45 minutes (**Fig. 4f**). While several CXC and CC chemokines were also differentially expressed in THP-1, their cognate receptors in K562 were poorly expressed, reducing the likelihood of these ligands driving the K562 NF-κB program.

We next evaluated whether prioritized ligands were induced within the putative causal window preceding K562 NF-κB activation, with TNF serving as an expected mediator (**Fig. 4g**). ChronoPack ranked *TNF* first, compared with ranks of 2, 3, 18-19 (tie), and 23-50 (tie) for NicheNet^29^, TraSig^30^, LIANA^31^, and CellChat^28^, respectively. ChronoPack generally favored ligands that showed strong, coordinated induction within this window (median log2FC: 1.8) whereas several ligands prioritized by other methods, including IL1B, exhibited weaker or delayed induction (0.01–0.9 median log2FC). Thus, by requiring ligand induction to precede downstream activation at a biologically plausible lag, ChronoPack identifies signaling interactions whose temporal dynamics are consistent with a causal role. More broadly, these findings demonstrate how dense temporal profiling can complement conventional ligand–receptor analyses by narrowing candidate signaling pathways based on their temporal compatibility.

### Stimulus-informed temporal dynamics improve gene regulatory network inference

We next investigated whether high time-resolution scRNA-seq could improve the inference of gene regulatory networks (GRNs). Inferring causal regulatory relationships from single-cell data remains challenging because many key regulators are activated post-translationally rather than transcriptionally. For example, TNF-α stimulation of K562 rapidly drives the nuclear translocation of NF-κB (NFKB1-RELA), whereas RELA transcript abundance remains largely unchanged. Consequently, co-expression-based methods may fail to recover regulatory relationships mediated by changes in TF activity that are not reflected in TF expression.

To address this limitation, ChronoPack integrates stimulus-associated signaling information with gene expression dynamics to prioritize regulatory interactions (**Fig. 5a**). Candidate TFs are ranked by their proximity to the stimulus in OmniPath signaling networks^32^, while candidate target genes are ranked by the significance of their temporal expression dynamics. These scores are combined to prioritize TF–target pairs, with the signaling component determining the relative priority of TFs and the expression component ranking candidate targets. Unlike co-expression-based methods, this approach does not require TF transcript abundance to reflect regulatory activity.

**Figure 5.**
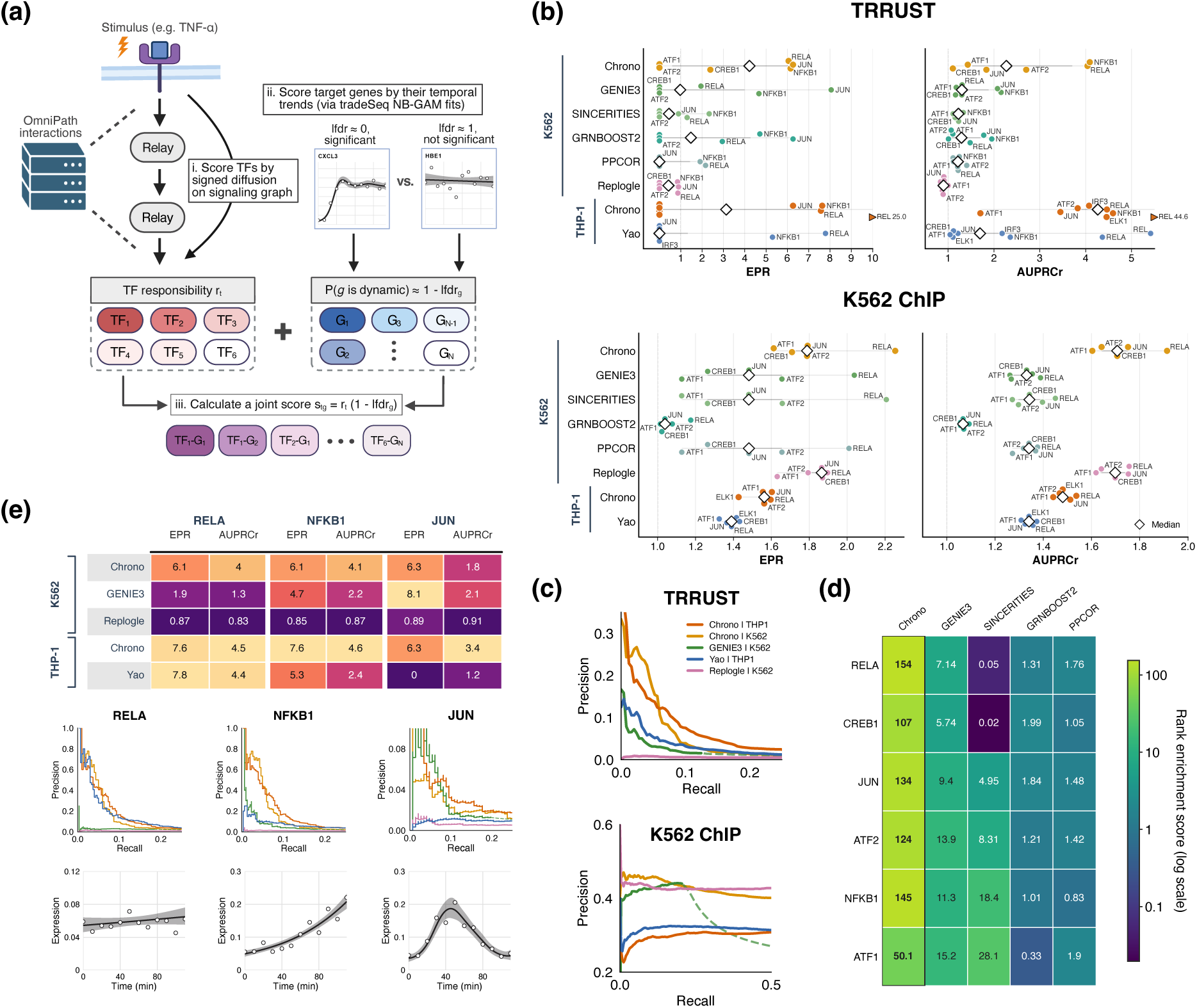
Stimulus-informed regulatory network prioritization with ChronoPack. **(a)** Overview of ChronoPack’s regulatory edge prioritization, which combines stimulus-proximal TF scores (calculated from diffusion on the signaling graph) with target gene dynamics. lfdr denotes the empirical local false discovery rate estimated from tradeSeq *associationTest* statistics, and is used to score genes by their evidence for dynamic expression. **(b)** Benchmarking of ChronoPack against co-expression-based methods and perturbation-derived rankings using the human TRRUST and K562 ChIP-seq reference networks. Performance is quantified using early precision ratio (EPR) and area-under-precision–recall-curve ratio (AUPRCr) (see definitions in **Methods**). All methods were evaluated against a common protein-coding gene universe, with unranked genes completed according to random expectation. TFs exceeding metric plot limits are denoted by triangles instead of circles and explicitly labeled with their exact values. **(c)** Macro-averaged precision-recall (PR) curves evaluated against the TRRUST and K562 ChIP reference networks. Dashed lines indicate expected precision after random completion of any unscored targets. **(d)** Rank enrichment of literature-supported inflammatory signaling TFs among predicted regulatory edges, reflecting their prioritization within each method’s inferred network. **(e)** TF-specific recovery of TRRUST interactions for RELA, NFKB1 and JUN. Heatmaps summarize EPR and AUPRCr, with corresponding PR curves and K562 expression trajectories shown for each TF.

We first evaluated target recovery for individual TFs against two complementary reference networks: curated human TRRUST interactions^33^ and promoter-proximal TF occupancy derived from K562 ChIP-seq. ChronoPack consistently outperformed established expression-based GRN inference methods^34–36^ in recovering TRRUST interactions in K562+TNF-α and achieved strong target recovery in THP-1+LPS (**Fig. 5b, c**). Across signaling-relevant TFs, ChronoPack achieved median early precision ratios (EPRs) of 3.1–4.2 and area-under-precision–recall-curve ratios (AUPRCr) of 2.3–4.3. By comparison, expression-based methods evaluated in K562 showed weaker and less consistent recovery (GRNBOOST2: median EPR = 1.5, AUPRCr = 1.3; GENIE3: median EPR = 1.0, AUPRCr = 1.3), particularly for TFs such as RELA. Against the K562 ChIP-seq reference, ChronoPack showed more modest enrichment (median EPR = 1.8, AUPRCr = 1.7), and its advantage over expression-based methods was largely explained by differences in candidate gene coverage. In contrast, its advantage against TRRUST persisted after controlling for coverage.

Perturbation-derived rankings (see **Methods**) revealed a contrasting pattern. The unstimulated K562 screen (Replogle et al.^37^) showed poor recovery of TRRUST interactions but performed comparably to ChronoPack against K562 ChIP-seq, whereas the LPS-stimulated THP-1 screen (Yao et al.^38^) achieved strong TRRUST recovery for several inflammatory regulators. These findings suggest that basal-state perturbation screens may incompletely capture regulatory programs engaged during acute inflammatory stimulation.

Consistent with its stimulus-informed prior, ChronoPack preferentially assigned regulatory influence to TFs with established roles in TNF-α or LPS signaling (**Fig. 5d**). Compared with expression-based methods, its predictions were substantially more enriched for signaling-relevant regulators such as RELA and JUN. The preferential recovery of signaling-relevant regulators and their known targets supports the complementary contributions of stimulus proximity and temporal gene expression dynamics to regulatory network inference.

To determine when stimulus-informed inference offers the greatest advantage, we compared target recovery for regulators with distinct routes of activation (**Fig. 5e**). For RELA, ChronoPack achieved strong target recovery in both K562 and THP-1 (EPR = 6.1–7.6). In K562, this substantially exceeded GENIE3 and Replogle-derived rankings (EPR = 1.9 and 0.87, respectively), whereas in THP-1, ChronoPack performed similarly to the stimulus-matched Yao-derived rankings (EPR = 7.8). Because RELA is activated primarily through signaling-dependent nuclear translocation rather than transcriptional induction, its transcript abundance provides limited information for expression-based GRN inference. By contrast, the induced regulators NFKB1 and JUN showed appreciable target recovery with GENIE3 (EPR = 4.7 and 8.1, respectively), compared with 6.1 and 6.3 for ChronoPack. GENIE3 slightly outperformed ChronoPack for JUN, consistent with expression-based approaches benefiting from variation in TF transcript levels. These comparisons highlight the particular advantage of stimulus-informed inference for regulators whose activity is poorly captured by transcript abundance, while demonstrating competitive performance without requiring TF–target-specific regulatory evidence.

### Inflammatory responses share a conserved repertoire of kinetic modes

To assess how transcriptional response kinetics may vary across cellular and signaling contexts, we compared gene dynamics across three inflammatory settings: direct TNF-α stimulation of K562 cells (K562 1°), secondary activation of K562 cells in co-culture with LPS-stimulated THP-1 cells (K562 2°), and direct LPS stimulation of THP-1 cells. Despite differences in stimulus source and cellular identity, genes repeatedly resolved into a limited set of dynamic modes, including transient induction, sustained activation, and repression (**Fig. 6a,b**). The relative abundance of these modes was notably similar across contexts, with impulse-up, trend-up, trend-down, and transient-down comprising 14-23%, 24-36%, 46-51%, and 3-6% of responsive genes, respectively. Gene membership, however, was considerably less conserved. While canonical NF-κB targets such as NFKBIA and CXCL3 retained broadly similar response kinetics across conditions, consistent with a shared NF-κB-centered inflammatory component, other genes were deployed more variably. HES1, for example, exhibited impulse-like induction in both K562 inflammatory settings but was repressed in LPS-stimulated THP-1 cells. Other genes, such as CCL3 and IRF1 (**Fig. 4c**), were only responsive and significantly dynamic in one cell type. Consistent with these observations, the two K562 conditions shared approximately 71% of top induced genes (see **Methods**), whereas overlap between K562 and THP-1 was substantially lower, at approximately 32-35%. Repressed genes were even less conserved overall, with only 12-36% shared across inflammatory contexts.

**Figure 6.**
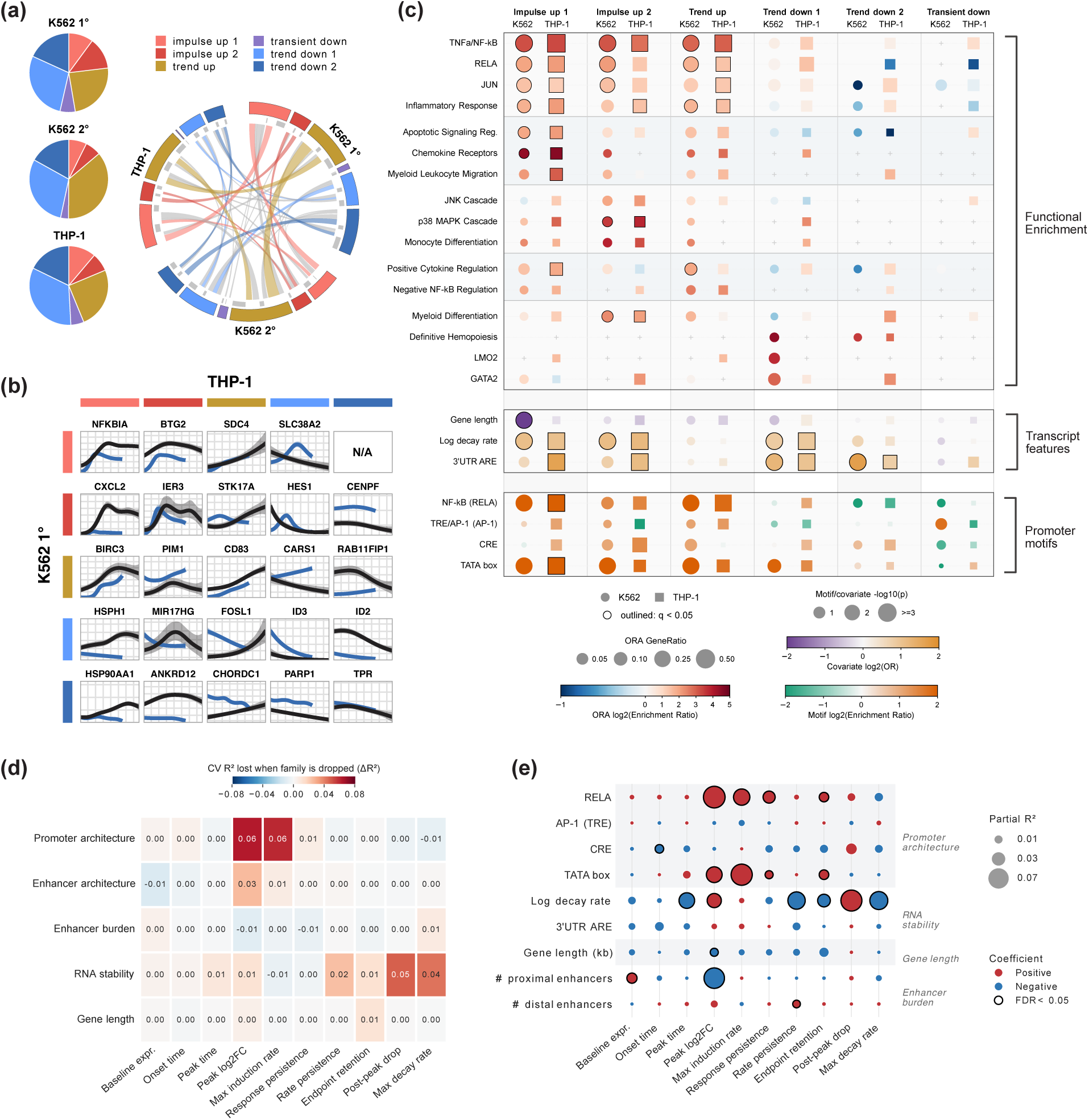
Inflammatory response modes are conserved but gene composition varies across cellular contexts. **(a)** Dynamic mode composition and reassignment of top dynamic genes across K562 1° (TNF-α stimulation), K562 2° (THP-1 co-culture), and THP-1 (LPS stimulation). **(b)** Representative genes with conserved or variable dynamics between K562 1° (blue) and THP-1 (black). **(c)** (*top panel*) Functional enrichment of dynamic modes. Induced genes are enriched for TNF/NF-κB, JUN/AP-1, cytokine, and inflammatory-associated signatures, whereas trend-down genes are enriched for lineage/state-associated programs. Dot size denotes GeneRatio, color denotes log2(enrichment ratio), shape denotes cell type, and black outlines indicate FDR < 0.05. (*middle and bottom panels*) Associations of transcript-level features and enrichment of transcription factor motifs in gene promoters. Induced modes are associated with NF-κB, AP-1/bZIP, and TATA box motifs in both K562 1° and THP-1. Dot size denotes −log10(p), color denotes log2(odds ratio) [transcript features] or log2(enrichment ratio) [motifs], shape denotes cell type, and black outlines indicate FDR < 0.05. **(d)** Unique predictive contribution of each feature family for different trajectory descriptors, calculated as the difference in held-out cross-validated R^2^ between the full elastic net model and a model excluding that family (ΔR^2^ = 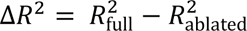). **(e)** Adjusted associations between selected regulatory features and trajectory descriptors from multivariable OLS models; dot size denotes partial R^2^, color indicates coefficient direction, and black outlines indicate FDR < 0.05. Associations are conditional on the other features included in the regression model.

ChronoSeq further revealed differences in the timing of these conserved response programs. Following alignment of the initial responses, induced genes in LPS-stimulated THP-1 cells exhibited slower onset than those in K562 cells, with impulse-up genes reaching peak expression substantially later (**Fig. S12**). These findings suggest that conserved transcriptional programs can nevertheless unfold on different timescales. This shift may partly reflect differences in receptor-proximal signal processing. TNF-α rapidly engages the TNFR–IKK–NF-κB pathway, whereas TLR4 partitions LPS signaling between an early MyD88-dependent arm and a delayed endosomal TRIF-dependent arm that sustains NF-κB activity and engages additional IRF-dependent programs.^39–41^ This signaling difference may contribute to the slower LPS response, although cell-type-specific regulatory architecture is also likely to contribute. Together, these results suggest that inflammatory responses preserve a broadly similar repertoire of kinetic programs, while their timing and gene membership can be extensively reshaped by cellular identity and signaling route. Importantly, dense temporal sampling provides a means to resolve these modes and characterize how response kinetics vary across cellular and signaling contexts.

### Conserved dynamic modes are associated with distinct regulatory architectures

The deployment of similar kinetic programs across diverse inflammatory contexts raises a fundamental question: to what extent do these shared response patterns reflect common regulatory architectures despite differences in cellular identity and signaling? To address this, we systematically examined how the dynamic modes differed in their transcript-level properties, cis-regulatory features, and associated functional programs.

Functional enrichment analysis revealed that the major dynamic modes mapped to distinct biological modules rather than reflecting arbitrary kinetic classes (**Fig. 6c**). Impulse-up 1 genes exhibited the strongest and most consistent enrichment for canonical TNF/NF-κB signaling in both K562 and THP-1 cells, including RELA and JUN regulons, inflammatory and apoptotic programs, and genes associated with chemokine signaling and leukocyte migration. By comparison, impulse-up 2 retained an inflammatory signature but showed greater enrichment for JNK and p38 MAPK-associated programs, particularly in THP-1 cells, indicating functional stratification even among transiently induced genes. Trend-up genes also remained enriched for NF-κB and AP-1 targets, but were comparatively associated with cytokine regulation and negative regulation of NF-κB, consistent with a more sustained regulatory and feedback component of the response. In contrast, trend-down genes showed little enrichment for canonical inflammatory programs and instead favored lineage-associated signatures, including definitive hematopoiesis and LMO2- and GATA2-associated programs in K562 and myeloid differentiation in THP-1. The smaller transient-down class showed comparatively little functional enrichment. Thus, the kinetic classes resolved the inflammatory response into distinct effector, signaling and regulatory programs while separating these from pre-existing lineage-associated programs.

The dynamic modes were further distinguished by two complementary regulatory layers: promoter architecture and transcript-level properties. NF-κB family motifs, represented by RELA, provided the clearest promoter signature of induction, showing broad enrichment across induced dynamic modes in both K562 and THP-1 cells, with particularly strong enrichment in impulse-up 1 and trend-up genes (**Fig. 6c**). TATA boxes were similarly favored among induced promoters, consistent with an initiation-ready core-promoter architecture previously associated with elevated basal TBP and promoter-proximal Pol II occupancy and rapid activation of immediate/early inflammatory genes.^42,43^ TRE/AP-1 motifs showed a weaker but predominantly induction-associated pattern, consistent with rapid MAPK-dependent activation of pre-existing AP-1 complexes, whereas CRE-like motifs were more context dependent, with a pronounced enrichment among THP-1 impulse-up 2 genes. These results suggest that inflammatory gene induction involves a broadly conserved NF-κB-associated promoter architecture, with additional regulatory motifs distinguishing kinetic modes and cellular contexts.

Beyond these promoter-level differences, transcript-level properties were more closely associated with overall response persistence. Both impulse-up classes were enriched for transcripts with higher estimated mRNA decay rates^44^ and 3′-UTR AU-rich elements^45^, whereas enrichment for these features was much weaker among trend-up genes, consistent with more sustained transcript accumulation. Shorter gene lengths were most evident among K562 impulse-up 1 genes and may further facilitate rapid transcriptional responses for a subset of immediate-response genes. Notably, turnover-associated features also extended to repressed genes: trend-down 1 genes showed elevated decay rates and ARE enrichment, while AREs remained prominent among trend-down 2 genes, suggesting that intrinsic transcript instability may likewise facilitate depletion as transcription declines. Thus, promoter architecture primarily distinguished inflammatory induction, whereas transcript stability more clearly separated transient from sustained induction and may additionally shape the rate of transcript loss during repression. Together, these findings suggest that conserved kinetic modes reflect shared combinations of functional programs, promoter architecture, and transcript-level properties across distinct inflammatory contexts.

### Promoter architecture and RNA turnover predict distinct dimensions of inflammatory gene expression kinetics

The distinct cis-regulatory and transcript-level features associated with the dynamic mode phenotypes suggested that different regulatory layers may shape separable dimensions of gene expression kinetics. To examine this directly, we decomposed induced trajectories into continuous measures of response magnitude, timing, persistence, and decay, and asked how well each kinetic descriptor could be predicted from gene-associated features. Using cross-validated elastic net models, promoter architecture carried the strongest unique predictive information for response magnitude: removing promoter features reduced held-out R^2^ by 0.064 for peak induction and 0.060 for the maximum induction rate (**Fig. 6d**).

Enhancer motifs provided a smaller additional contribution to peak induction (ΔR^2^ = 0.026), with this signal arising primarily from distal enhancer-like elements. In contrast, RNA stability features were much more informative than motif architecture for later response dynamics, particularly post-peak decline (ΔR² = 0.048), maximum observed decay rate (ΔR² = 0.044), and other measures of persistence and attenuation. Enhancer CRE burden and transcript length carried little unique predictive information across these trajectory dimensions, while onset and peak timing were largely unpredictable from the promoter, enhancer, and transcript features examined here. Overall, full-model predictive performance was modest across trajectory descriptors (cross-validated R² ≈ 0-0.29; **Fig. S13**). This separation is consistent with previous work showing that core-promoter composition can influence transcriptional output and inducibility, whereas RNA turnover can strongly shape the temporal progression of inflammatory gene expression.^46–49^

Feature-level associations further resolved the specific features underlying this separation. RELA motif incidence and TATA box architecture were associated with greater response magnitude (peak log2FC, maximum induction rate), with RELA additionally associated with greater persistence after the response peak (**Fig. 6e**). By contrast, mRNA decay rate was most strongly associated with post-peak decline, reduced expression persistence, endpoint retention, and maximum observed decay, while AU-rich elements showed broadly concordant but weaker associations across these kinetic dimensions. Together, these results support a modular view of inflammatory gene regulation, in which promoter motif architecture is most informative of induction strength, while transcript-intrinsic turnover is more closely linked to response duration. The ability to resolve these relationships across continuous trajectory dimensions illustrates how dense temporal profiling can move beyond descriptive kinetic classes to identify distinct regulatory layers associated with different phases of a response. Such a framework should enable dynamic gene expression phenotypes to be linked more directly to regulatory mechanisms and provide a basis for investigating how transcriptional control and RNA turnover shape response trajectories.

## Discussion

Single-cell genomics has largely been built around static snapshots. While this framework has transformed our ability to catalog cellular states, it provides only indirect access to the dynamic processes that generate those states. Many regulatory events, including signal transduction, transcriptional activation, transcript decay, and intercellular communication, unfold on timescales of minutes, yet conventional scRNA-seq workflows remain constrained by manual sampling, processing latency, and sparse temporal coverage. ChronoSeq and ChronoPack were developed to address this limitation by enabling automated acquisition and analysis of densely sampled single-cell transcriptional trajectories. Rather than treating transcription as a static cellular property, this framework allows transcriptional responses to be studied as dynamic processes evolving through time.

The value of dense temporal sampling was immediately evident in the investigation of both intracellular and intercellular response programs. In TNF-α-stimulated K562 cells, ChronoSeq revealed a pre-existing erythroid-like subpopulation exhibiting substantially attenuated NF-κB responses relative to the remaining K562 cells. This finding illustrates how baseline cellular heterogeneity can shape subsequent response dynamics and demonstrates the value of characterizing these differences over time rather than relying on static endpoint measurements. Similarly, in THP-1–K562 co-culture, dense temporal profiling captured the delayed propagation of inflammatory signaling from LPS-stimulated macrophages to neighboring K562 cells, with TNF emerging as a leading candidate mediator. Temporal ordering, therefore, may provide an orthogonal source of information for studying multicellular communication, complementing existing approaches based on static ligand–receptor co-expression.

Building on previous studies of early inflammatory transcriptional dynamics^50,51^, ChronoSeq enabled the systematic characterization of genome-wide response kinetics at minute-scale single-cell resolution. The repertoire of kinetic modes was broadly conserved across contexts, whereas as their gene composition and timing varied substantially. These modes were associated with distinct biological functions and regulatory features. Further analysis of individual kinetic parameters revealed how different molecular properties relate to specific aspects of gene transcription: promoter motif architecture was most predictive of induction magnitude, enhancer motifs provided a smaller additional contribution, and transcript stability was more closely associated with post-peak attenuation and response duration. Induced genes shared a common inflammatory promoter architecture centered on NF-κB, while differences in transcript stability and other transcript-level properties were more closely related to whether responses were transient or sustained. Repressed genes, by contrast, were preferentially associated with lineage-linked programs, pointing to coordinated attenuation of pre-existing cellular states during activation. Together, these findings point to a conserved regulatory logic in which promoter architecture helps define stimulus responsiveness, while transcript-level properties shape the duration and form of the resulting expression response.

Dense temporal sampling also improves the inference of stimulus-responsive gene regulatory networks. Many central signaling transcription factors, including RELA, are activated primarily through post-translational mechanisms and therefore leave little information in TF–target mRNA covariance. By integrating stimulus context with the temporal dynamics of the transcriptional response, ChronoPack recovered known regulatory interactions more effectively than conventional co-expression methods and performed competitively with perturbation-derived rankings for stimulus-responsive transcription factors. Interestingly, this performance was achieved without requiring TF–gene-specific evidence, indicating that signaling context and temporal response dynamics can themselves provide a strong prior for network inference. These findings highlight the value of temporal information as a complementary source of evidence for regulatory inference, particularly when transcription factor activity is decoupled from its own mRNA abundance.

Despite its robust performance, we acknowledge certain limitations of the current ChronoSeq platform. First, while our system captures temporal dynamics at unprecedented resolution, it still requires cell lysis; therefore, trajectories are computationally reconstructed from dense populations rather than reflecting true longitudinal monitoring of the exact same physical cell (as in Live-seq). Second, the current hardware iteration is limited to a 12-plex Time-Tag barcode system. While 12 timepoints are sufficient for acute, fast-acting responses (0–3 hours), mapping slower processes like multi-day differentiation will require engineering larger Time-Tag libraries. Finally, as with many high-throughput microfluidic methods, the per-cell transcript recovery is lower than standard commercial snapshot platforms. However, comparisons with matched 10X Chromium data recovered the expected transcriptional programs, and the reduced per-cell transcript yield had minimal impact on temporal trajectory reconstruction and characterization.

Looking forward, we anticipate that high-density temporal datasets will become increasingly important for building predictive models of cellular behavior. Just as large collections of static single-cell profiles enabled systematic characterization of cellular states, densely sampled temporal trajectories may enable quantitative models that predict how those states emerge, transition, and respond to perturbation. By providing a practical framework for measuring transcriptional dynamics at minute-scale resolution, ChronoSeq opens a path toward a more mechanistic and quantitative understanding of cellular decision-making.

## Data Availability

ChronoSeq data are available in the ArrayExpress database (http://www.ebi.ac.uk/arrayexpress) under accession numbers E-MTAB-15894, E-MTAB-15928, E-MTAB-15927, E-MTAB-15946, E-MTAB-15947, E-MTAB-15956, E-MTAB-15955, E-MTAB-15964, E-MTAB-15965 and E-MTAB-15966. Parameters used for each ChronoSeq experiment and the associated accession numbers are listed in **Table 1**.

**Table 1:** Experimental parameters used for various ChronoSeq datasets and their accession numbers.

| **Experimental condition** | **Cell concentration** | **Time-Tags** | **Stimulus** | **Sampling interval** | **Stimulation time** | **EMBL-EBI ArrayExpress accession number** |
|---|---|---|---|---|---|---|
| K562 | 430 cells/µl | 12 | 10ng/ml TNF-α | 10 minutes | 5 minutes after first sample | E-MTAB-15894 |
| 50:50 K562 + differentiated THP-1 cells | 430 cells/µl | 12 | 1µg/ml LPS | 15 minutes | 5 minutes after first sample | E-MTAB-15928 |
| K562 | 500 cells/µl | 12 | 10ng/ml TNF-α + 5µg/ml Puromycin | 10 minutes | Both added together 5 minutes after first sample | E-MTAB-15947 |
| K562 Negative Control (Reverse Time-Tag injection order 12 to 1) (QC) | 860 cells/µl | 12 | No Stimulus. Time-Tag Bias estimation | 10 minutes | N/A | E-MTAB-15946 |
| 50:50 K562+EL4 (QC) | 215 cells/ µl | 3 | 10ng/ml TNF-α | 10 minutes | 5 minutes after first sample | E-MTAB-15955 |
| 50:50 K562+EL4 (QC) | 430 cells/ µl | 3 | 10ng/ml TNF-α | 10 minutes | 5 minutes after first sample | E-MTAB-15956 |
| 50:50 U937+K562 | 430 cells/ µl | 12 | 0.5µg/ml LPS | 10 minutes | 5 minutes after first sample | E-MTAB-15927 |
| K562+EL4 (Time Tag 10), K562(Time Tag 11), EL4 (Time Tag 12) (QC) | 215 cells/ µl | 3 | No Stimulus. Only Checking Time-Tags | 7 minutes | N/A | E-MTAB-15964 |
| Bulk K562 and EL4 cells mixed with 12 Time-Tags for verification (QC) | 600 cells/ µl | 12 | No Stimulus. Only Checking Time-Tags | N/A | N/A | E-MTAB-15965 |
| Bulk K562 cells negative control (QC) | 600 cells/ µl | 12 | No Stimulus | 10 minutes | N/A | E-MTAB-15966 |

## Code Availability

ChronoPack code, documentation, tutorials, and figures are available at GitHub: https://github.com/Wang-lab-UCSD/ChronoPack.

## Online Methods

### Experimental procedures

Detailed experimental procedures and protocols can be found on the ChronoSeq project website http://wanglab.ucsd.edu/ChronoSeq/. The notebooks associated with the various protocols also feature embedded YouTube videos. See **Supplementary Table S2** for items purchased for assembling the ChronoSeq device. For the latest and most accurate links and information always refer to the protocols and instructions on the project website.

### Device design overview

Our device design (**Figure S8**) addresses several challenges not present in the original Drop-seq^52^ system. See **Figure S9** for a comparison and detailed discussion of the ChronoSeq and Drop-seq devices. The system distinguishes itself from conventional Drop-seq devices through several key innovations: automated bead handling with vortex suspension, integrated cell culture system with 37°C water bath and 5% CO₂ environment, robotic droplet collection and bead recovery system, and compressed air pressure control with flow sensing for rapid flow rate changes.

### Microfluidic chip design and fabrication

See **Figure S10** for a comparison of the ChronoSeq and Drop-seq chip designs and the reasons for the various design changes. For fabrication standard photolithography and PDMS molding techniques^53^ are used for chip fabrication. Chips are plasma bonded to glass slides and chemically treated to create hydrophobic channels. We implemented a rigorous quality control protocol: compressed air testing to check for delamination and eliminate leaking chips, simulating the high pressures used during the flushing phase; visual inspection to ensure the absence of particles before connecting to the device. These design and quality control measures enable the chips to withstand higher pressures and cyclical pressurization during the flushing phase. The optimized chip design is crucial for reliable, high-throughput, time-resolved single-cell sequencing. The improvements in channel geometry and manufacturing process contribute to reduced clogging, increased cell capture efficiency, and overall system robustness. These features are essential for maintaining consistent performance across multiple time points and ensuring high-quality data collection throughout extended experimental runs.

### Creation of Time-Tagged ChronoSeq beads from Drop-seq beads

ChronoSeq beads were prepared by modifying Drop-seq beads (**Supplementary Figure S3**). The modification principle involves binding oligonucleotides (SEQ1 through SEQ12 **Supplementary Table S3**) to the existing polyT region of Drop-seq beads, followed by extension with E. coli DNA Polymerase I to create the reverse complement of the oligo, and subsequent alkaline denaturation to generate single-stranded DNA. Drop-seq beads were obtained from LGC Biosearch Technologies (14-nt UMI). Exact protocol used can be found in **Supplementary Protocol 1.**

### Library preparation

ChronoSeq uses the same oligos used for Drop-seq library preparation (**Supplementary Table S3).** ChronoSeq library preparation protocol (**Supplementary Protocol 2**) has an expected yield between 1.5-6ng/ μL after the first PCR reaction as measured by Qubit. While Tagmented sequencing-ready libraries have an expected concentrations of 20-50 ng/μL. Libraries were sequenced on one lane of Illumina NovaSeq X Plus 10B with 10% PhiX Spike-In and Custom Read 1 Primer mixed with Standard Illumina Primers. Either PE100 or PE150 configurations were used for the sequencing run. The ChronoSeq protocol is very similar to the Drop-Seq V3.1 protocol with the following notable changes. First, the Exonuclease 1 treatment step was omitted since it did not contribute to Library yield, quality or capture rates. Next, 14 PCR cycles were typically done instead of 12 PCR Cycles recommended by Drop-Seq for the first round of PCR. Finally, beads were split into different reactions based on number of Timepoints instead of number of estimated STAMPs for both PCR and Reverse Transcription.

### Cell culture

All cell lines were cultured in DMEM supplemented with 10% fetal bovine serum (FBS) and 1% penicillin-streptomycin (Pen-Strep) at 37°C in 5% CO₂. THP-1 cells were additionally differentiated into macrophages with 100 nM phorbol 12-myristate 13-acetate (PMA) for 48 hours, then rested for 72 hours. THP-1 macrophages were then converted to a cell suspension non-enzymatically with EDTA to preserve cell surface proteins. **Supplementary Protocol 3** provides exact details of cell culture conditions for each cell line.

### qPCR validation

K562 cells were sampled with the ChronoSeq device at 10-minute intervals over 2 hours, yielding 12 timepoints. Following baseline sampling, TNF-α (10 ng/mL) was introduced at the 5-minute mark to induce inflammatory signaling. Cell lysates were prepared using a modified Bio-rad SingleShot Cell Lysis Kit protocol (**Supplementary Protocol 4)** in which the cell suspension was directly mixed with the Lysis reagents instead of washing and changing the media to DPBS.

### qPCR validation of K562 co-stimulation by THP-1 cells and detection of RNase activity

To investigate paracrine signaling effects between immune cell populations, THP-1 cells were differentiated and stimulated with 1 μg/ml LPS, with cell supernatants collected at 0, 1, 2, and 3 hours post-stimulation. These time-course supernatants were then applied to K562 target cells. Controls included unstimulated K562 cells and K562 cells stimulated with TNF-α. K562 cells were sampled at 0 and 1 hour following supernatant addition using a modified Bio-rad SingleShot Cell Lysis Kit protocol (**Supplementary Protocol 5).** Next, we performed qPCR for the relatively lowly expressed housekeeping gene HPRT1 in some of the same samples of K562 cells stimulated with THP-1 supernatants and controls (**Fig. S7**). HPRT1 was not detected in all samples stimulated with THP-1 supernatants, with no Cq values generated (See **Table S1**). Based on the same data we also observed no RNase activity for K562 cells stimulated with TNF-α. This confirmed high RNase activity is a problem unique to THP-1 cell suspensions. Thus, we added 3U/µl of Watchmaker Genomics RNase inhibitor (from a stock concentration of 800U/µl) for each Time-Tag in our THP-1-K562 mixture experiment.

### Computational procedures

#### ChronoSeq-Tools pipeline

The ChronoSeq-Tools pipeline extends the Drop-seq pipeline methodologies to enable temporal tracking of single-cell transcriptomes through integrated Time-Tags. The pipeline processes paired-end RNA sequencing data through sequential barcode extraction, alignment, and Time-Tag determination phases to generate a final Digital Gene Expression matrix (DGE) with Time-Tags for each Cell. Scripts and instructions for running the ChronoSeq-Tools pipeline can be found on GitHub https://github.com/kanishkasthana/ChronoSeq-Tools. Moreover, code for generating QC plots in this paper can be found at https://github.com/kanishkasthana/ChronoSeq-QC

### Pipeline architecture

The workflow initiates by converting FASTQ files to unaligned BAM format using Picard’s FastqToSam. Three sequential tagging steps employ Drop-seq’s TagBamWithReadSequenceExtended to isolate:

1. **Cell barcodes** (XC tag: bases 1-12)
2. **Unique molecular identifiers** (XM tag: bases 13-26)
3. **Time-Tags** (YT tag: bases 27-86) with no base quality thresholds.

Quality control steps include filtering reads with low-quality tags and trimming adapter sequences (AAGCAGTGGTATCAACGCAGAGTGAATGGG) and polyA tails using Drop-seq tools. Processed reads undergo STAR alignment against species-specific references, followed by queryname sorting and MergeBamAlignment to preserve metadata. Gene annotation occurs via TagReadWithGeneFunction using RefFlat files, generating digital expression matrices(DGEs). Computational efficiency is enhanced through optional scratch directory usage and distributed locking mechanisms to manage I/O operations on SLURM cluster systems.

#### Alignment references

For species-mixing experiments reads were aligned to the same hg19-mm10 mixed cell reference used in Dropseq. For all other experiments hg38 was used for alignment.

### Time-Tag determination

Time tags derive from twelve synthesized oligonucleotides containing a combination of degenerate bases (R,Y,M,K,S,W,H,B,V,D,N) and constant bases flanked by polyA sequences. The GetTimeTags algorithm:

1. Extracts YT tag sequences from aligned BAM files
2. Performs reverse complement conversion
3. Matches sequences against regex patterns for each Time-Tag.
4. Requires ≥20 detections per cell barcode for reliable assignment.
5. Assigns final Time-Tags through majority vote (≥70% prevalence) for each Cell barcode. Cell barcodes failing to meet this threshold are flagged as having “Time Tag Collision Detected” indicating potential cross-contamination or sequencing errors

### Comparison with Drop-seq pipeline

While sharing core components (Picard, Drop-seq tools, STAR alignment), ChronoSeq introduces three key modifications:

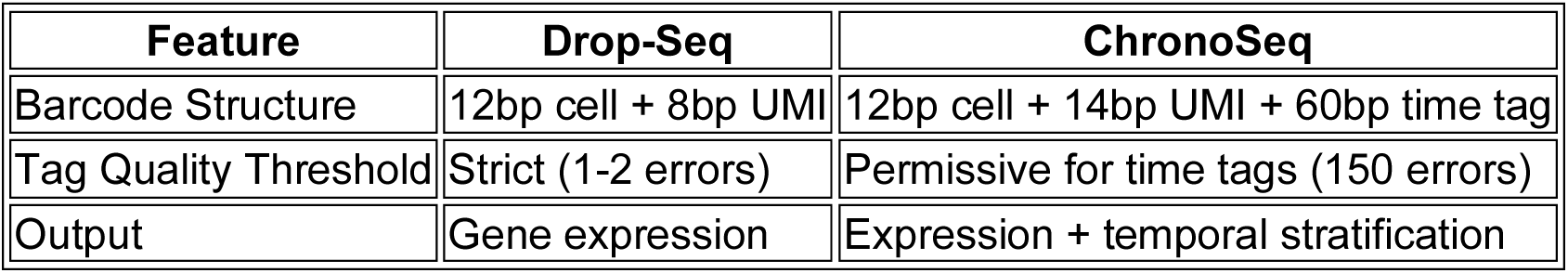

| Feature | Drop-Seq | ChronoSeq |
|---|---|---|
| Barcode Structure | 12bp cell + 8bp UMI | 12bp cell + 14bp UMI + 60bp time tag |
| Tag Quality Threshold | Strict (1-2 errors) | Permissive for time tags (150 errors) |
| Output | Gene expression | Expression + temporal stratification |

### Optional error correction

ChronoSeq supports optional error correction for both substitution and synthesis errors in cell barcodes, closely mirroring Drop-seq’s established methodology. Our substitution error correction has been optimized to run on multiple cores to speed up execution. Like Drop-seq the correction step identifies cell barcodes differing by a single base (Hamming distance of 1) and collapses those with a dominant sequence representing at least 93% of reads, mitigating sequencing errors. The synthesis error correction step identifies and collapses barcodes with systematic errors at the barcode termini, typically arising from bead synthesis imperfections. This is accomplished by constructing a Prefix-Trie data structure and merging barcodes that differ only at the last few positions, replacing the error-prone bases with ‘N’ and updating associated UMIs accordingly. Using a Prefix-Trie approach allows us to speed up execution from Quadratic to Linear time for our approach compared to Drop-Seq. These corrections are performed prior to digital gene expression quantification and time tag assignment, ensuring high-fidelity cell barcode recovery.

### ChronoPack package

#### Standard preprocessing

Time-Tagged ChronoSeq digital gene expression (DGE) matrices generated using the ChronoSeq-Tools pipeline were processed using Scanpy^54^. Cells with fewer than 500 detected genes or 500 UMIs, genes detected in fewer than three cells, and cells identified as doublets using Scrublet^55^ were removed. Additional cell quality filters were applied to total UMI counts (>15,000 for K562+TNF-α and >10,000 for THP-1/K562+LPS) and mitochondrial transcript fractions (>5% and >7.5%, respectively). Scrublet was applied separately to each reverse-transcription batch, and an additional low-quality batch was excluded from the co-culture experiment. Following quality control, UMI counts were normalized to 10,000 per cell and log-transformed (*sc.pp.normalize_total*, *sc.pp.log1p*). Cell-cycle scores were calculated using published S- and G2/M-phase marker genes^56^.

For K562+TNF-α, the top 3,000 highly variable genes were selected across reverse-transcription batches. Total UMI counts, mitochondrial transcript fraction, and cell-cycle scores were regressed out, followed by ComBat^57^ correction for reverse-transcription batch effects. Additional low-quality cell clusters were excluded following Leiden clustering. For THP-1/K562+LPS, the top 2,000 highly variable genes were selected after excluding cell-cycle genes, and total UMI counts were regressed out without additional batch correction. The processed expression matrices were scaled and subjected to principal component analysis (PCA), followed by neighborhood graph construction, UMAP visualization, and Leiden clustering. K562 and THP-1 cells in the co-culture experiment were distinguished using lineage-specific marker genes.

#### Deriving a transition kernel from optimal transport couplings

WaddingtonOT^20^ was used to infer transport couplings between cells at successive timepoints using highly variable gene expression. For K562+TNF-α, couplings were computed from batch-corrected expression, whereas K562 and THP-1 cells in the co-culture experiment were analyzed separately using log-normalized expression. Default parameters for the entropy, source marginal, and target marginal regularization terms were used (ɛ = 0.05, λ_1_ = 1, λ_2_ = 50), with three iterations for growth-rate estimation. The resultant couplings were stitched together into a sparse transport matrix *T* with *T_ij_* encoding the descendant mass sent from cell *i* to cell *j*. For each cell, only the 30 largest forward couplings were retained, yielding a pruned transport matrix *T*. The forward transition kernel *P* was obtained by row-normalizing this matrix, such that *P* = *D*^−1^*T* where *D* = diag(*T*1), and *P_ij_* is the probability of cell *i* transitioning to cell *j*. For the co-culture experiment, the resulting K562 and THP-1 transition kernels were assembled into a block-diagonal matrix. *P* was used directly for several downstream tasks, including cell-wise trajectory clustering and cell fate mapping.

#### Cell-wise trajectory clustering

We used the transition kernel *P* to cluster cells according to their inferred future trajectories. For each cell, we estimated a probability distribution over individual terminal cells by propagating its transition probabilities to the final measured time point. Specifically, for a cell *i* observed at time point *r*, we calculated *S_ij_* = (*P^N^*^−*r*^)*_ij_*, where *N* denotes the final sampling index and *S_ij_* denotes the probability that cell *i* reaches terminal cell *j* after *N* − *r* transition steps. Thus, each row of *S* represents the inferred distribution of terminal destinations for a given cell. We then used cosine similarity between these terminal-cell distributions to construct a weighted nearest-neighbor graph, such that cells with similar predicted futures were assigned stronger pairwise similarities. The graph was constructed using the 500 most similar neighbors per cell, symmetrized, and pruned to a maximum of 500 connections per cell. We then applied Leiden clustering (sc.tl.leiden; resolution = 0.3 for K562+TNF-α and 0.6 for the co-culture experiment) to identify trajectory clusters, defined as communities of cells with shared future-state distributions.

#### Cell fate mapping

Terminal fates were defined by the trajectory cluster assignments of cells at the final sampled time point. To estimate fate probabilities, we used the WaddingtonOT transition kernel *P*, which connects cells at consecutive time points through strictly forward-directed transitions. Because the resulting graph is acyclic, fate probabilities can be calculated efficiently through backward recursion rather than by solving the linear system of a general absorbing Markov chain. For each fate *A_F_*, terminal cells were assigned a probability of 1 if they belonged to *A_F_* and 0 if they belong to another terminal fate. Starting from these terminal cells, we propagated the probabilities of reaching individual terminal cells backward through successive time points and summed them within each fate. Equivalently, fate probabilities satisfy the recursion *p_F_*(*i*) = ∑*_j_ P_ij_* · *p_F_*(*j*), where *P_ij_* denotes the probability of transitioning from cell *i* to cell *j*. Fate uncertainty was quantified using normalized Shannon entropy *H*(*i*) = − ∑*_F_ p_F_*(*i*) · log *p_F_*(*i*)⁄log *K*, where *K* = 2 is the number of terminal fates.

To identify early transcriptional features associated with the inferred response trajectories, we correlated gene expression at the first sampled time point with each cell’s fate contrast score, *y_i_* = *p*_1_(*i*) − *p*_0_(*i*). Using log-normalized expression values, Spearman rank correlations were calculated after excluding genes detected in fewer than 10% of early cells. P-values were adjusted for multiple testing using the Benjamini–Hochberg procedure, and significant positive and negative correlations (*q* < 0.05) were associated with fates *A*_1_ and *A*_0_, respectively.

#### Technical gene filtering

To identify genes exhibiting technical variation associated with individual Time-Tags, we compared time-tag-resolved pseudobulk expression profiles from each experiment against an unstimulated K562 bulk negative-control dataset processed with the same Time-Tag barcodes. UMI counts were aggregated across cells within each Time-Tag, normalized to counts per million (CPM), and log-transformed as log(1 + CPM). For each gene present in both datasets, expression profiles across the 12 corresponding Time-Tags were independently smoothed using cubic splines (SciPy UnivariateSpline, *k* = 3), with smoothing parameter s = 0.5 × n × Var(y) where *n* denotes the number of timepoints and Var(*y*) the variance of the corresponding expression profile. Residuals were calculated by subtracting the smoothed expression profiles from the observed values. We then calculated the Pearson correlation (r) between experimental and negative-control residuals and the fraction of experimental expression variance associated with control-correlated residual variation, *f* = max(*r*^2^, 0) × Var(residuals)⁄Var(expression). Genes with *r* ≥ 0.7 and *f* ≥ 0.25 were excluded as candidates for time-tag-associated technical bias. This filtering was applied independently to K562 cells from the TNF-α experiment and to K562 and THP-1 cells from the LPS experiment, using the same K562 negative-control reference.

Following time-tag bias filtering, genes with mitochondrial (MT-) or ribosomal protein (RPL/RPS) prefixes, genes lacking annotated gene symbols, and genes with fewer than 10 total UMIs across the corresponding cell population were excluded before tradeSeq fitting. The resulting gene sets were used for temporal modeling and downstream analyses of gene expression dynamics.

#### Identifying genes with significant temporal trends

For each gene, we used the tradeSeq package^24^ to fit a negative-binomial generalized additive model (NB-GAM), modeling gene expression as a smooth function of sampling time. Gene-level UMI counts were modeled as *Y_gi_* ∼ NB(µ_gi_, ϕ*_g_*) with log µ_gi_ = log *N_i_* + *α_g_* + ∑*_l_ Z_li_s_gl_*(*t_i_*) where *Y_gi_* denotes the count for gene g in cell i, *N_i_* is the cell-specific library size normalization factor, *α_g_* is a gene-specific intercept, *Z_li_* indicates the lineage assignment of cell *i*, and *s_gl_*(*t_i_*) is a lineage-specific smooth function of time. Each smoother was represented using cubic regression splines, *s_gl_*(*t_i_*) = ∑*_k_* b*_k_*(t*_i_*)β*_glk_*.

Model fitting proceeded in two stages. First, NB-GAMs with four spline knots were fitted to all genes that passed quality filtering, using a relatively constrained spline basis to reduce sensitivity to noise during genome-wide testing. Temporal variation in expression was evaluated using tradeSeq’s *associationTest*, which applies a Wald test to evaluate the null hypothesis that mean gene expression remains constant throughout the observed time course. P-values were adjusted across genes using the Benjamini-Hochberg procedure, with nominally significant genes (*p* < 0.05) retained for downstream analysis. These genes were subsequently refitted using eight spline knots to better capture rapid and transient changes in expression. All tradeSeq association-test statistics and *p*-values were derived from the initial four-knot fits. For analyses requiring high-confidence dynamic genes, we additionally required FDR-adjusted *q* < 0.05 and |peak log2FC| > 0.5.

### Quantification of individual gene expression trajectories

Trajectory features were calculated from fitted gene expression profiles y(t) evaluated on a common time grid resampled to one-minute intervals. Fitted values were represented on the log(1 + *x*) scale. For each gene, expression at the first sampled time point was designated as the baseline.

#### Response amplitude and peak timing

To accommodate both induced and repressed responses, the response peak was defined as the time point with the largest absolute deviation from baseline: *t_peak_* = argmax*_t_*|*y*(*t*) − *y*(*t*_0_)|, where *y*(*t*_0_) denotes expression at the first time point. Ties were resolved in favor of the earliest time point. We recorded baseline expression, peak expression, and peak change via their difference: Δ*_peak_* = *y*(*t_peak_*) − *y*(*t*_0_). Peak fold change was additionally calculated after back-transforming the log-expression values: peak log2FC = log_2_ ((*e^y^*^(*t*^*^peak^*^)^ − 1)⁄(*e^y^*^(*t*0)^ − 1)).

#### Response onset time

Onset was estimated by identifying sustained deviation from an early baseline reference. Each gene trajectory was first standardized across time to have mean 0 and standard deviation 1. The reference was then defined as the mean standardized expression observed during the first 10 minutes and was treated as constant across the full observation window. At each time point, deviation from this early reference was calculated as Δ*_z_* = *z*(*t*) − mean(*z*(*t* ≤ 10min)). The onset time was defined as the first time point at which |Δ*_z_*| > 0.20 for at least thirty minutes. The absolute deviation criterion enabled detection of both upward and downward responses. Genes that did not satisfy this criterion during the observation window were assigned a missing onset time.

#### Normalized trajectory area

To summarize the position of each trajectory within its observed expression range, expression values were independently rescaled to the interval from 0 to 1: *y*_01_(*t*) = (*y*(*t*) − min(*y*))⁄(max(*y*) − min(*y*)). The area under the rescaled trajectory was calculated and divided by the total duration of the observation window: AUC_norm_ = AUC*_y_*_01(*t*)_⁄(*t_max_* − *t_min_*). This metric ranges from 0 to 1 and represents the fraction of the observed interval for which expression remains near the upper portion of the gene’s own dynamic range. Constant trajectories were assigned an AUC_norm_ value of 0.

#### Trajectory shape

Shape features were calculated after independently rescaling time and expression to the interval from 0 to 1, allowing trajectory geometry to be compared independently of response magnitude and experiment duration. At each time point, curvature was estimated as: 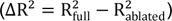, where *y*^′^ rescaled and *y*^′′^_rescaled_ denote finite-difference first and second derivatives with respect to the normalized time axis. Larger values indicate stronger local curvature in the trajectory. We retained the mean and maximum curvature for each gene, representing overall bending and the strongest localized change in shape, respectively. As a complementary measure, we calculated the turning angle between consecutive trajectory segments within the rescaled trajectories. At each time point, the angle between the incoming and outgoing segments was calculated from their vector dot product and divided by *π*. The normalized turning angle ranged from 0 to 1, with values near 0 indicating locally straight trajectories and larger values indicating sharper changes in direction. For each gene, we retained the mean and maximum normalized turning angles, representing overall directional variability and the sharpest local turn, respectively.

#### Response and rate persistence

For each fitted curve, expression was first baseline adjusted by subtracting the fitted value at the initial time point and truncating negative values to zero: *y*(*t*) = max(*y*(*t*) − *y*(*t*_0_), 0). To quantify how well induced expression was maintained after reaching its maximum, we computed the running maximum envelope *m*(*t*) = max_τ≤*t*_ *y*(τ) and integrated both *y*(*t*) and *m*(*t*). Response persistence was defined as P = AUC_*y*_⁄AUC*_m_* such that values near 1 indicate monotonic or sustained induction, whereas lower values indicate transient responses that decline after an earlier peak. Rate persistence was calculated analogously by using the finite-difference derivative of the expression curve.

#### Induction and decay rates

Rates of expression change were estimated between consecutive points of the fitted trajectory as *v_i_* = (*y*(*t_i_*_+1_) − *y*(*t_i_*))/(*t_i_*_+1_ − *t_i_*), with each rate assigned to the midpoint of the corresponding time interval. The maximum induction rate was defined as the largest rate observed at or before the response peak, *v*_ind_ = max*_ti_*≤ *t*_peak_ *v_i_*. The maximum decay rate was defined as the most negative rate observed at or after the response peak, *v*_decay_ = min*_ti_* ≥ *t*_peak_ *v_i_*, such that more negative values indicate faster post-peak attenuation.

#### Post-peak attenuation and endpoint retention

The absolute loss of expression following the response peak was quantified as Δ_post-peak_ = y(t_peak_) − y(t_end_) where t_end_ denotes the final time point. For induced responses, larger positive values therefore indicate greater attenuation after the peak. Endpoint retention quantified the fraction of the peak response remaining at the end of the observation window: *R*_end_ = (*y*(*t*_end_) − *y*(*t*_0_))⁄(*y*(*t*_peak_) − *y*(*t*_0_)). A value of 1 indicates complete retention of the peak response, a value of 0 indicates a return to baseline, and negative values indicate expression below the initial baseline at the endpoint.

### Synthetic downsampling analysis

To assess the effect of temporal sampling density on gene trajectory reconstruction, we synthetically downsampled the K562+TNF-α time course from 10-minute to 30-minute intervals. For a matched comparison, both datasets were restricted to the first 90 minutes, retaining all cells at each selected timepoint (5,674 cells across 10 timepoints for the full-resolution dataset and 2,582 cells across four timepoints for the downsampled dataset). NB-GAMs were fitted independently using eight spline knots for the full-resolution data, consistent with the standard ChronoPack pipeline, and four knots for the downsampled data to accommodate its reduced number of unique timepoints. Onset and peak times were compared between the full-resolution and downsampled fits for 238 high-confidence induced genes (*q* < 0.05, |peak log2FC| > 0.5).

Peak times were evaluated for all 238 genes, while onset comparisons were restricted to the 229 genes with defined onset times in both fits. Estimation errors were calculated as the absolute deviation from the matched full-resolution fit and summarized as mean ± standard error. Representative gene trajectories were additionally compared to evaluate changes in reconstructed response shape.

### Clustering genes into dynamic response modes

To classify temporal response shapes, we extracted tradeSeq-fitted trajectories for genes with nominally significant dynamics (*p* < 0.05; K562+TNF-α: 2,022; K562+THP-1+LPS: 1,920 K562 and 1,119 THP-1 genes) and aligned them to a shared one-minute grid using population-specific time offsets and scaling factors. These parameters were selected by visually aligning the initial transcriptional responses across experiments, with additional temporal scaling of the THP-1 response to approximately match the onset-to-peak interval of rapidly induced NF-κB targets, including NFKBIA, in stimulated K562 cells. Trajectories were restricted to the common aligned time window and row-normalized to emphasize temporal shape rather than expression magnitude. To capture both overall trajectory shape and the timing of expression changes, each gene was represented by its normalized expression trajectory together with its first temporal derivative. These components were weighted such that expression and derivative information contributed 90% and 10%, respectively, to the squared Euclidean distance between genes.

Induced and repressed genes, defined by the sign of their peak log2 fold change, were clustered separately by agglomerative hierarchical clustering using Ward’s minimum-variance criterion (Ward.D2). For each response direction, we retained two nested resolutions of the Ward dendrogram. Major dynamic modes were selected across *K* = 2-100 by jointly considering mean silhouette width and resampling stability. Stability was assessed across 200 80%-gene subsamples using the adjusted Rand index (ARI) for overall partition agreement and Jaccard similarity for individual cluster recovery, with these metrics supporting K=3 for both induced and repressed genes. This yielded three dynamic modes in each direction: impulse-up 1, impulse-up 2 and trend-up for induced genes, and trend-down 1, trend-down 2 and transient-down for repressed genes. To identify finer trajectory structure within the major modes, we retained a second, higher-resolution cut of the same dendrogram based on the within-cluster sum-of-squares (WSS) elbow across *K* = 2-100. This yielded 17 induced and 16 repressed subclusters overall, with 4–7 assigned to each dynamic mode.

### Gene conservation across dynamic modes

To quantify how consistently genes retained the same dynamic mode across inflammatory contexts, we calculated a directional recovery score. For each mode, high-confidence dynamic genes (*q* < 0.05, |peak log2FC| > 0.5) in the source context were ranked by their NB-GAM Wald statistic, and the top *k* = 100 genes were retained. Directional recovery was defined as the fraction of these genes that remained high-confidence and retained the same mode assignment in the target context. This approach prioritized the strongest mode-associated genes in the source context while allowing their recovery among all high-confidence genes in the target context, irrespective of rank. This calculation was performed in both directions, and the mean of the two directional recovery fractions was used as the final mode conservation score.

### Cell–cell communication

#### ChronoPack

Candidate ligand–receptor (L-R) interactions between THP-1 sender cells and K562 receiver cells were evaluated using the secreted signaling subset of the human CellChatDB database^28^ Interactions were retained when all ligand genes were detected in THP-1 cells and all receptor genes were detected in K562 cells, defined as nonzero counts in more than 5% of cells in the corresponding population. For multi-component ligand or receptor complexes, annotations were decomposed into their constituent gene symbols, and all required subunits were required to satisfy the same detection criterion. This expression-based filtering reduced the CellChatDB secreted signaling catalog from 1,199 to 230 candidate interactions.

Receptor complexes were summarized by the geometric mean of their constituent subunits. For each retained L-R pair, we computed a time-resolved activity score, *A_lr_*(*t*) = √*L_l_*(*t*) · *R̅_r_*, where *L_l_*(*t*) denotes ligand expression in THP-1 cells at time *t*, and *R̅_r_* is the mean expression of receptor *r* in K562 cells across the time course. Activity trajectories were baseline-subtracted relative to the first measured time point, and each L-R pair was summarized by its summed activity over an early causal window, defined to be at least 30 minutes prior to NF-κB response onset in K562. Candidate interactions were then prioritized by z-score normalizing L–R activity and ligand temporal significance, represented as − log_10_(*p*), and summing those two quantities into a joint score.

#### LIANA and CellChat

were evaluated using a time-agnostic analysis in which cells were pooled across post-stimulation time points into THP-1 sender and K562 receiver populations. Cells from the initial time point were excluded to avoid combining unstimulated and LPS-stimulated THP-1 states. LIANA rank aggregation was performed on the log-transformed expression layer using the consensus ligand–receptor resource, with ligands and receptors required to be expressed in at least 10% of cells and each group required to contain at least five cells. CellChat was implemented through the LIANA wrapper.

#### TraSig

Experimental time was scaled to the interval [0,1] and used as pseudotime, with THP-1 and K562 cells assigned to sender and receiver trajectories, respectively. Multi-subunit L-R complexes were excluded because TraSig does not provide built-in aggregation of complex expression. Inference was performed using default settings (--metric dot, --startingTreatment smallerWindow, --alignType unaligned, and --nLap 20, --numPerms 10000). Interactions were ranked by dot-product score, and both empirical and false-discovery-rate-adjusted P values were retained.

#### NicheNet

Candidate ligands were restricted to those detected in THP-1 cells before the onset of the K562 response (0–75 min). Receiver targets were defined as K562 genes classified by ChronoPack as both significantly dynamic (P < 0.05) and induced (peak log2 fold change > 0.5), with all detected genes used as the background. Using the NicheNet v2 human ligand–target prior, each ligand was scored according to how well its weighted target predictions ranked receiver target genes above background. Ligands were then prioritized by corrected area under the precision–recall curve, calculated as AUPRC minus target prevalence in the background.

ChronoPack was benchmarked against LIANA rank aggregation, CellChat, TraSig, and NicheNet. The candidate interaction universe comprised human CellChatDB interactions annotated as secreted signaling, restricted to pairs for which the ligand components were detected in THP-1 cells and the receptor components were detected in K562 cells. Receptor-resolved methods were evaluated using complete ligand–receptor interaction identities, whereas NicheNet was evaluated using ligand identity. Positive-control recovery was quantified as the best rank assigned to TNF–TNFRSF1A, TNF–TNFRSF1B, or TNF. To assess whether each method prioritized ligands induced before the K562 response, ligand induction was quantified over a putative causal window restricted to time points at least 30 min before response onset. For each ligand, the change in expression across this window was calculated as the log2 fold change. Method performance was summarized as the median log2 fold change among the ten highest-ranked ligands.

### Gene regulatory network inference and benchmarking

TF–gene regulatory edges were prioritized by combining independent evidence for stimulus-proximal regulators and responsive target genes. For each gene, temporal evidence was derived from the tradeSeq association test. Association test *P* values were modeled using a beta-uniform mixture to estimate a local false discovery rate (lfdr*_g_*), with target gene scores defined as *s_g_* = 1 − lfdr*_g_*. This quantity was used as an empirical non-null score, such that higher values indicated stronger evidence that gene *g* exhibits dynamic expression over time.

Regulator priors were derived from OmniPath^32^ post-translational signaling interactions. We constructed a directed, signed signaling network in which stimulatory and inhibitory interactions were represented by positive and negative edges, respectively; self-interactions were removed, compound nodes were expanded to individual genes, and duplicate interactions were consolidated while retaining sign information where available. Signed absorbing diffusion was initiated from the stimulus, with activating and inhibitory signal propagated separately, inhibitory edges reversing signal polarity, and transcription factors treated as absorbing terminal nodes. A leakage term progressively penalized longer signaling paths. For each TF, absorbed signaling evidence was summarized as total support, defined as the sum of absorbed positive and negative mass, and net activation, defined as absorbed positive minus absorbed negative mass. The latter quantity provided a TF-level measure of stimulus-dependent activation; positive values were retained and normalized across candidate TFs to define the regulator score *r_t_*.

Candidate TF-gene edges were then assigned the joint score *s_tg_* = *r_t_*(1 − lfdr*_g_*), thereby prioritizing edges for which both the regulator was supported by the upstream signaling context and the target showed strong temporal evidence of regulation. Self-edges were excluded. Because regulator and target evidence were estimated independently, the edge score did not incorporate TF–gene-specific interaction information.

ChronoPack was benchmarked against four expression-based GRN inference methods implemented through BEELINE^58^ – GRNBoost2^36^, GENIE3^59^, PPCOR^35^ and SINCERITIES^34^ – as well as TF-gene rankings derived from the genome-scale K562 Perturb-seq dataset of Replogle *et al.*^37^ and the 3-h LPS-stimulated THP-1 Perturb-seq dataset of Yao *et al.*^38^ Processed single-cell data for the Replogle and Yao datasets were obtained from the authors’ Figshare repository and GEO submission (GSM6858447), respectively. Differential expression was calculated independently for each TF perturbation using Seurat^60^ FindMarkers, comparing cells carrying a targeting guide with non-targeting controls. Each perturbed TF was treated as a regulator and differentially expressed genes as candidate targets, with edges ranked by − log_10_(*P*).

Networks were evaluated against curated human TRRUST^33^ regulatory interactions and a K562 ChIP-seq reference network. Analyses focused on stimulus-relevant TFs represented in each reference. The core set comprised RELA, NFKB1, JUN, CREB1, ATF1 and ATF2, with ELK1, REL and IRF3 additionally considered in the THP-1 context. Reference-specific availability further restricted the evaluated sets: ATF2 was absent from the Yao dataset, NFKB1 was not represented in the K562 ChIP reference, and REL and IRF3 were not included in the ChIP-based comparisons.

The K562 ChIP reference was constructed from ENCODE K562 ChIP-seq peak calls as a binary TF–gene promoter-overlap network. Released GRCh38 narrowPeak files were obtained for factors annotated as transcription factors, prioritizing optimal IDR-thresholded peak sets and using conservative IDR sets when optimal peaks were unavailable. One representative peak set was retained per TF, and peaks were filtered to signalValue ≥ 10. Promoters were defined from GENCODE v45 basic protein-coding annotations as strand-aware intervals extending 1 kb upstream and 200 bp downstream of the transcription start site, with one promoter retained per gene. Peaks were intersected with promoter intervals using BEDTools, and a TF–gene edge was assigned when at least one peak overlapped the corresponding promoter. This reference therefore captures promoter-proximal TF occupancy in ENCODE K562 cells rather than direct evidence of functional transcriptional regulation.

All benchmarks were evaluated over a common universe of 19,338 protein-coding genes, with self-edges removed. Performance was quantified using early precision ratio (EPR) and area-under-the-precision–recall-curve ratio (AUPRCr). EPR measures enrichment of true positives among the top-*k* ranked edges, where *k* is the number of reference targets for the evaluated TF, relative to random expectation. AUPRCr was calculated as the area under the precision–recall curve divided by the expected AUPRC of a random ranking, given by the prevalence of reference-positive edges. EPR therefore emphasizes recovery among the highest-ranked predictions, whereas AUPRCr summarizes performance across the full TF-specific network. These metrics were quantified separately for each TF against its corresponding reference targets, with aggregate performance summarized by the median across TFs and macro-averaged precision-recall curves.

To evaluate regulator-level prioritization independently of target recovery, we performed a rank-enrichment analysis analogous to an unweighted GSEA running-sum statistic. For each method, TF–gene edges were ranked by their method-specific scores after removing duplicate TF–target pairs and self-edges. For each TF, edges assigned to that regulator were treated as hits and all remaining edges as misses, and the enrichment score (ES) was defined as the maximum positive difference between the cumulative fractions of hits and misses along the ranked edge list. The rank at which this maximum occurred was recorded as the peak-enrichment rank. To account for differences in the number of edges assigned to each TF, null distributions were generated by randomly redistributing the same number of TF-specific edges across the ranked list. Normalized enrichment scores (NES) were calculated as the observed ES divided by the mean maximum ES under this matched null, and empirical *P* values were calculated as (1 + *n_null_*_≥*obs*_)/(1 + *B*), where *B* is the number of permutations.

### Cell-type-specific mapping of cis-regulatory elements to genes

For K562, cell-type-resolved candidate cis-regulatory element (cCRE) annotations were obtained from ENCODE file ENCFF286VQG, which provides K562-specific promoter-like signature (PLS), proximal enhancer-like signature (pELS), and distal enhancer-like signature (dELS) classifications from the ENCODE Registry of cCREs v4 on the GRCh38 genome assembly. Gene-proximal PLS and pELS elements were assigned to protein-coding genes by intersecting their genomic coordinates with ±200bp and ±2kb windows, respectively, around transcription start sites defined by GENCODE v45. Distal enhancer–gene links were obtained from ENCODE-rE2G^61^ predictions in ENCFF269DKY, which integrate chromatin accessibility, histone modifications, and chromatin contact information to nominate target genes for distal regulatory elements.

Because precomputed THP-1-specific cCRE annotations were unavailable, active THP-1 cCREs were defined by intersecting the genome-wide ENCODE v4 cCRE registry with THP-1 chromatin accessibility data. DNase-seq peaks from ENCODE experiment ENCSR896ADX, peak file ENCFF359IAQ, were used as a proxy for accessible chromatin, and cCREs overlapping a DNase-seq peak were retained as active. This procedure identified 71,912 accessible cCREs, comprising 17,450 PLS, 26,266 pELS, and 28,196 dELS elements. As in K562, PLS and pELS elements were assigned to genes using ±200bp and ±2kb windows around GENCODE v45 transcription start sites. Because rE2G predictions were not available for THP-1, distal enhancer–gene associations were instead defined using EPIraction^62^ enhancer–promoter interaction predictions from ENCFF848NJE.

### Gene-level feature analysis

To identify gene-level features associated with the distinct transcriptional response modes, we performed cell-type-stratified univariate logistic regression analyses in K562+TNF-α and THP-1+LPS. Each of the six dynamic modes — impulse-up 1, impulse-up 2, trend-up, transient-down, trend-down 1 or trend-down 2 — was modeled separately against the remaining genes in the corresponding cell-type-specific gene universe retained after technical filtering. The background universes contained 14,342 genes for K562 and 14,645 genes for THP-1. We evaluated 12 gene-level features comprising gene length, transcript decay rate, 3′-UTR AU-rich element (ARE) status, and features derived from three classes of linked candidate cis-regulatory elements (cCREs): promoter-like (PLS), proximal enhancer-like (pELS) and distal enhancer-like (dELS) elements. For each cCRE class, we quantified the number of linked elements, their presence or absence, and their aggregate GC content. Continuous predictors were z-score standardized within each cell type and foreground-background comparison, whereas binary predictors were retained as indicator variables. Missing values were excluded on a model-specific basis without imputation. For cCRE GC-content analyses, models were restricted to genes linked to at least one element of the corresponding cCRE class.

#### Gene length

Gene length was defined as the genomic span from transcription start to transcript end, including introns. A representative transcript was selected using a hierarchical preference for MANE-, CCDS-, and GENCODE basic-annotated protein-coding transcripts, with the median transcript span used within the highest-priority annotation class available for each gene.

#### Transcript decay rate

Transcript stability measurements were obtained from the Transcriptome Turnover Database (TTDB)^44^, which aggregates high-throughput RNA-turnover measurements across published studies. Reported decay rates were log10-transformed and summarized for each gene by their median value.

#### AU-rich element status

ARE annotations were obtained from ARED-Plus^45^, a curated resource of human 3′-UTR AU-rich elements. Transcript-level annotations were collapsed to the gene level, and genes were classified as ARE-positive when at least one annotated transcript contained a 3′UTR ARE.

#### GC content

For each gene and cCRE class, GC content was calculated as %GC = ∑*_i_*(*G_i_* + *C_i_*)⁄∑*_i_ L_i_*, where *G_i_* + *C_i_* is the number of *G* and *C* bases of linked element *i*, and ∑*_i_ L_i_* is the total interval length of the linked elements.

Each predictor was tested independently using logistic regression implemented in the Python statsmodels package^63^, yielding 144 models across two cell types, six dynamic modes and 12 features. Effect sizes were reported as odds ratios; for continuous predictors, estimates represent the change in odds of membership in the focal dynamic mode per one-standard-deviation increase in the predictor, whereas binary-variable estimates represent presence versus absence. Statistical significance was assessed using two-sided Wald tests, with Benjamini–Hochberg correction applied across all 144 tests and separately within each cell type.

### Gene set enrichment across dynamic response modes

Over-representation analysis (ORA) was performed for 12 foreground gene sets corresponding to six dynamic modes (impulse-up 1, impulse-up 2, trend-up, trend-down 1, trend-down 2 and transient-down) in the K562+TNF-α and THP-1+LPS experiments. For each cell-type–mode combination, high-confidence dynamic genes (*q* < 0.05, |peak log2FC| > 0.5) were ranked by their tradeSeq association-test Wald statistic, and up to the top 100 genes were retained, yielding foreground sets of 35–100 genes each. These foreground sets were also used for subsequent motif enrichment and gene trajectory feature analyses. Cell-type-specific background universes comprised genes retained after technical filtering (14,342 genes for K562 and 14,645 genes for THP-1). Gene set collections included Hallmark, Gene Ontology Biological Process, Reactome and KEGG Legacy pathways obtained using msigdbr (MSigDB 2026.1)^64,65^, together with CollecTRI^66^ transcription factor regulons and seven curated gene sets representing immediate-early responses and monocyte/macrophage or leukocyte differentiation programs. Standard collections were tested using clusterProfiler::enricher()^67^, with gene sets restricted to 5–5,000 genes after intersection with the corresponding background universe; curated gene sets were tested using the equivalent hypergeometric test implemented in scipy.stats.hypergeom. Fold enrichment was calculated as the ratio of the foreground and background gene proportions. For the standard collections, Benjamini–Hochberg correction was applied jointly across all tests within each database; for curated gene sets, correction was performed separately for each gene set across the 12 foregrounds. Adjusted *P* < 0.05 was considered significant.

### Motif enrichment in gene-linked cis-regulatory elements

Enrichment of known transcription factor motifs was assessed in candidate cis-regulatory elements (cCREs) linked to the same 12 cell-type–mode foreground gene sets used for gene set enrichment analysis, representing six dynamic response modes in the K562+TNF-α and THP-1+LPS experiments. These sets contained 35–100 high-confidence genes per mode, as defined above. Genes were linked to promoter-like (PLS), proximal enhancer-like (pELS) and distal enhancer-like (dELS) elements using cell-type-specific gene–cCRE maps.

To limit disproportionate contributions from genes linked to many regulatory elements, the number of cCREs retained per gene was capped at the cell type- and cCRE class-specific median. These caps were one PLS per gene in both cell types, three and two pELSs in K562 and THP-1, respectively, and six and eight dELSs, respectively. When a gene exceeded the corresponding cap, linked elements were randomly subsampled in five independent replicates using fixed seeds, and intervals shared by multiple genes were collapsed by genomic coordinates.

Motif enrichment was evaluated separately for each cell type, dynamic mode, cCRE class and sampling replicate using HOMER^68^ findMotifsGenome.pl against the hg38 reference genome, preserving the native widths of the input intervals (-size given) and disabling de novo motif discovery (-nomotif). Rather than using randomly sampled genomic regions, each foreground was compared with a cCRE-matched background comprising all unique elements of the same cCRE class in the corresponding cell type, excluding foreground intervals from that replicate. Known motifs were tested separately using the JASPAR 2026 CORE vertebrate non-redundant collection^69^ and the vertebrate known-motif library distributed with HOMER. For the TATA-Box motif in PLSs, enrichment was additionally evaluated using a strict positional filter that counted only motif hits whose centers fell 35–25 bp upstream of the associated transcription start site, corresponding to the canonical TATA-box window.

For each motif, foreground and background hit counts were compared using a two-sided Fisher’s exact test. Benjamini–Hochberg correction was applied within each cell type, dynamic mode, cCRE class, motif database and sampling replicate. Effect size was reported as the signed log2 odds ratio, calculated with a 0.5 pseudocount to accommodate zero-count cells.

Reproducibility across subsampling replicates was summarized by requiring concordant effect direction across replicates and recording motifs that reached FDR < 0.05 in at least three of five replicates.

To reduce redundancy among closely related motifs, the combined JASPAR and HOMER libraries (1,490 motifs) were grouped into 204 sequence-similarity archetypes using a procedure adapted from Vierstra et al.^70^ Motif position-weight matrices were trimmed to their informative cores by removing flanking positions with information content below 0.25 bits, while retaining a minimum motif width where possible. Pairwise motif similarities, including reverse-complement comparisons, were calculated using TOMTOM/tomtom-lite^71,72^, symmetrized and converted to distances for complete-linkage hierarchical clustering. Cutting the resulting dendrogram at a distance threshold of 0.85 yielded 204 motif archetypes. When multiple closely related motifs showed similar enrichment patterns, representative motifs were selected from these archetypes for visualization.

### Prediction of continuous gene trajectory features

To determine whether distinct regulatory layers predict different dimensions of gene-expression kinetics, we analyzed genes from the three induced-mode foreground sets (impulse-up 1, impulse-up 2 and trend-up) defined above for ORA and motif enrichment, in the K562+TNF-α and THP-1+LPS experiments. Of the 559 gene–cell-type observations in these foreground sets, 534 had corresponding regulatory feature annotations and were retained for analysis, representing 484 unique genes. Fitted trajectories were summarized by ten continuous descriptors spanning baseline expression, response timing, magnitude, persistence and decay: baseline expression, onset time, peak time, peak log2 fold change, maximum induction rate, response persistence, rate persistence, endpoint retention, post-peak drop and maximum observed decay rate. Genes peaking at the end of the observation window were assigned a maximum observed decay rate of zero, indicating that no decline was observed within the sampled interval.

Predictors were organized into complementary regulatory feature classes. *Promoter architecture* included promoter motif incidence for 66 JASPAR and HOMER motifs drawn from four PWM-similarity archetypes dominated by AP-1/CRE-bZIP and NF-κB/Rel motifs, together with TATA box status, promoter number and span in kilobases, and cell type-matched promoter accessibility. *Enhancer architecture* comprised the corresponding 66 motif-incidence features within cell type-matched proximal (pELS) and distal (dELS) CREs. *Enhancer CRE burden* captured the number and total span of linked proximal and distal CREs. *RNA stability* comprised median mRNA decay rate and AU-rich element status, and *gene length* was represented by gene span. Cell type was included as an adjustment term in pooled models. Trajectory descriptors, dynamic mode assignments, association statistics and identifiers were excluded from the predictor set.

Predictive performance was evaluated using repeated nested elastic net regression, with genes grouped across folds so that observations corresponding to the same gene in different cell types were never split between training and test sets. Five outer and five inner folds were used, and the analysis was repeated ten times with matched outer-fold assignments across model specifications. Feature filtering, imputation, standardization and hyperparameter selection were performed using training data only. Held-out predictions from all outer folds were concatenated within each repeat to calculate *R*^2^ and mean absolute error. Models containing only cell type were evaluated using the same cross-validation splits to establish baseline predictive performance (**Fig. S13**). To quantify unique predictive information, feature families were removed individually or in predefined combinations from the full model, with contribution defined as 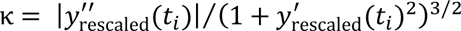. Family-only models were additionally used to measure the total predictive information contained within each feature class.

Individual feature associations were assessed separately using a multivariable ordinary least-squares model containing nine gene-associated predictors: RELA, AP-1 (TRE) and CRE promoter motif indicators, TATA box status, median mRNA decay rate, 3’UTR AU-rich element status, gene length, and pELS and dELS counts. Cell type was included as an additional adjustment covariate. Trajectory descriptors and continuous predictors were standardized, whereas binary predictors remained unscaled. Statistical significance was assessed using robust standard errors clustered by gene to account for genes represented in both cell types, with 95% confidence intervals estimated from 1,000 gene-level bootstrap resamples. Effect sizes were summarized by standardized regression coefficients and partial *R*^2^, and Benjamini–Hochberg correction was applied across the 90 predictor–outcome tests (9 predictors × 10 trajectory descriptors).

## Supplementary Figures and Tables

**Supplementary Figure S1.**
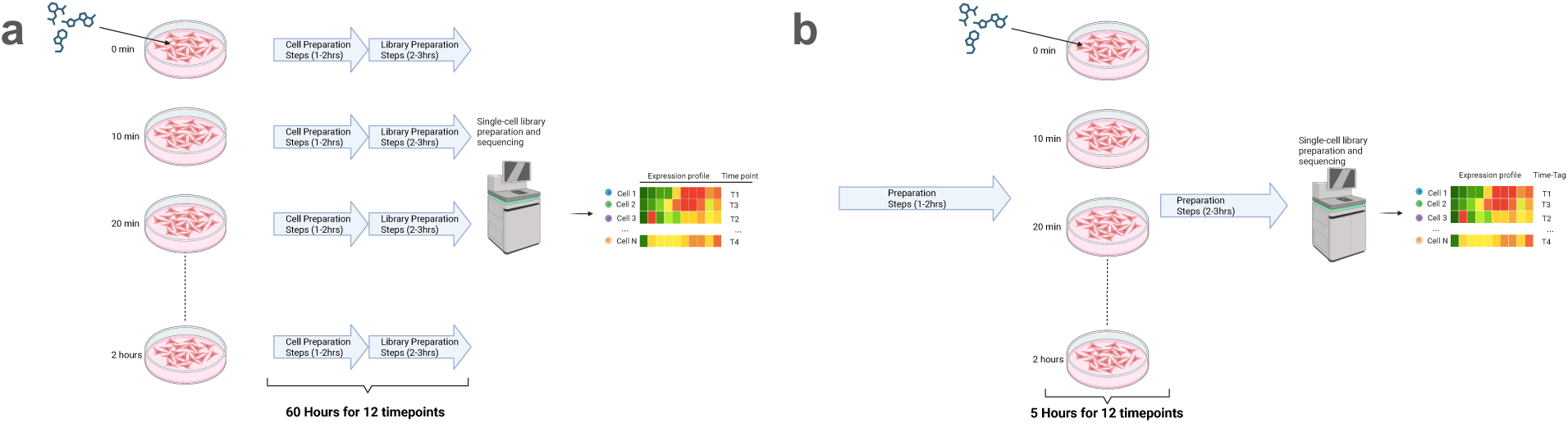
a) Shows the current paradigm for a temporal drug response study. If twelve timepoints are taken over two hours, each timepoint would have to be processed independently. Optimistically if five hours of hands-on labor is needed for each timepoint, this would require a minimum of 60 hours of total hands-on labor for all timepoints. Due to the lack of automation and manual nature of the technology there is a high possibility of inconsistent sampling or labelling times for each time point. For example, in each sample the metabolites used for labelling need to be washed out. Washing out is not instantaneous and the cells have time to incorporate more metabolites during processing delays; thereby increasing the chance of introducing an error of several minutes or longer for each sample. An error of several minutes introduces the high possibility of measuring spurious perturbations, especially if the sampling rate is also several minutes. Therefore, the high labor requirement, lack of automation, separate labeling of each sample, no immediate cell lysis to preserve transcriptome and resulting minute-scale sampling/labelling errors, make it practically impossible to use metabolic labelling for profiling real-time perturbations over several hours. b) If this same temporal drug response study is done using ChronoSeq this would require only five hours of hands-on labor. Moreover, by automating sample collection across time points and consolidating them into a single library via time-barcoded beads, ChronoSeq achieves minute-level sampling of expression dynamics over multiple hours. Automation and the immediate cell lysis help eliminate human errors in sampling and labelling, while integrated cell culture helps maintain optimal cell viability and function for experiments.

**Supplementary Figure S2:**
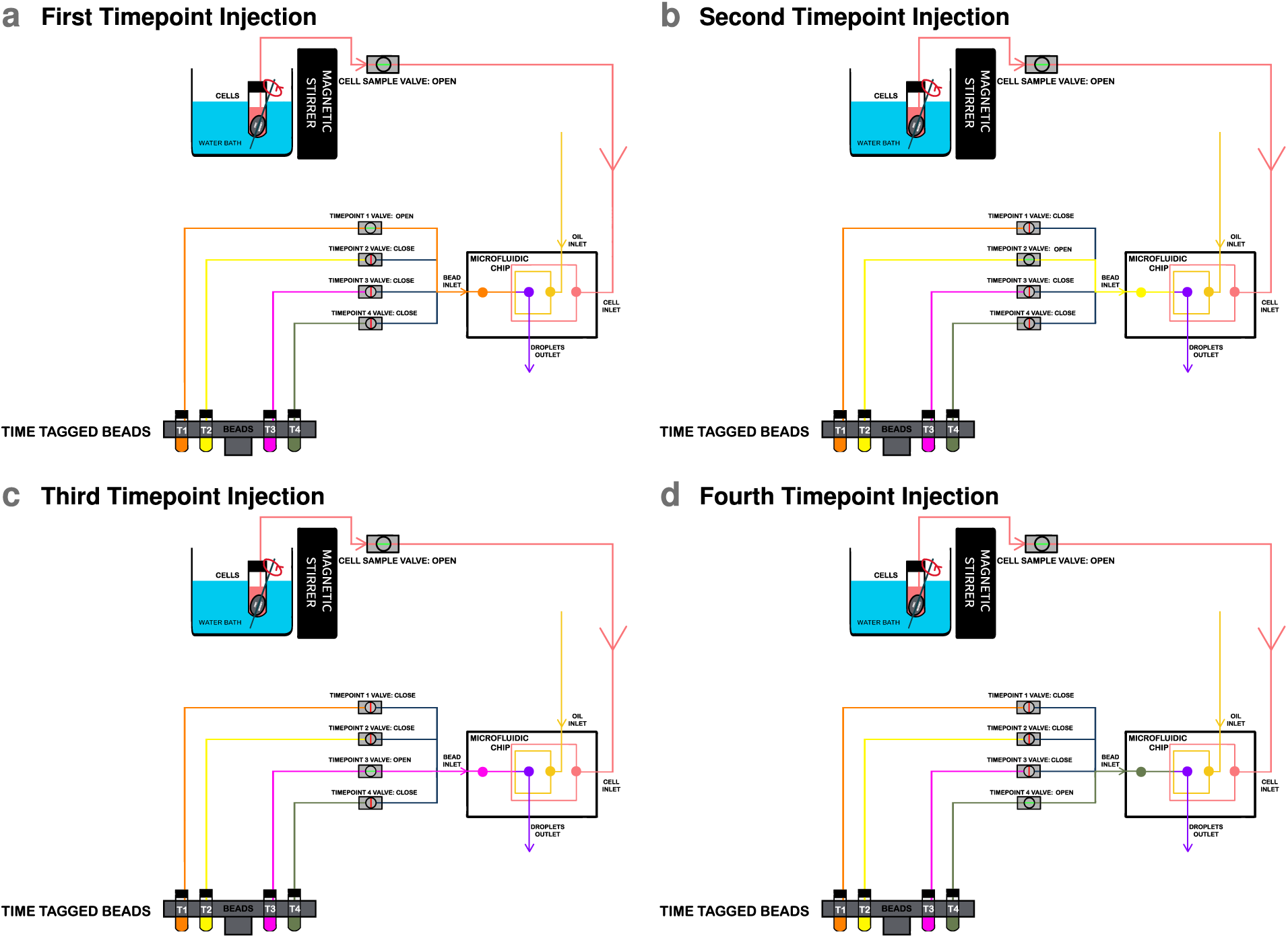
Simplified schematics to explain temporal barcoding using ChronoSeq device. The device sequentially co-injects Time-Tagged beads and cells into a droplet-generating microfluidic chip at regular time intervals. Panels a–d show the first to fourth timepoint injections, respectively. These beads are stored in separate reservoirs (e.g., 4 reservoirs for 4 time points for illustration purpose), with each reservoir containing beads with a unique Time-Tag. For each timepoint, the Cell Sample Valve remains open while the corresponding Timepoint Valve (orange for T1, yellow for T2, Hot Pink for T3, and Olive Green for T4) is opened. This allows for the co-injection of time-specific beads with cells, resulting in libraries barcoded with distinct Time-Tags. This process enables precise temporal labeling of cells across different timepoints. Our system currently supports 12 unique Time-Tags with minimum 7-minute intervals including a 1-minute sampling duration.

**Supplementary Figure S3:**
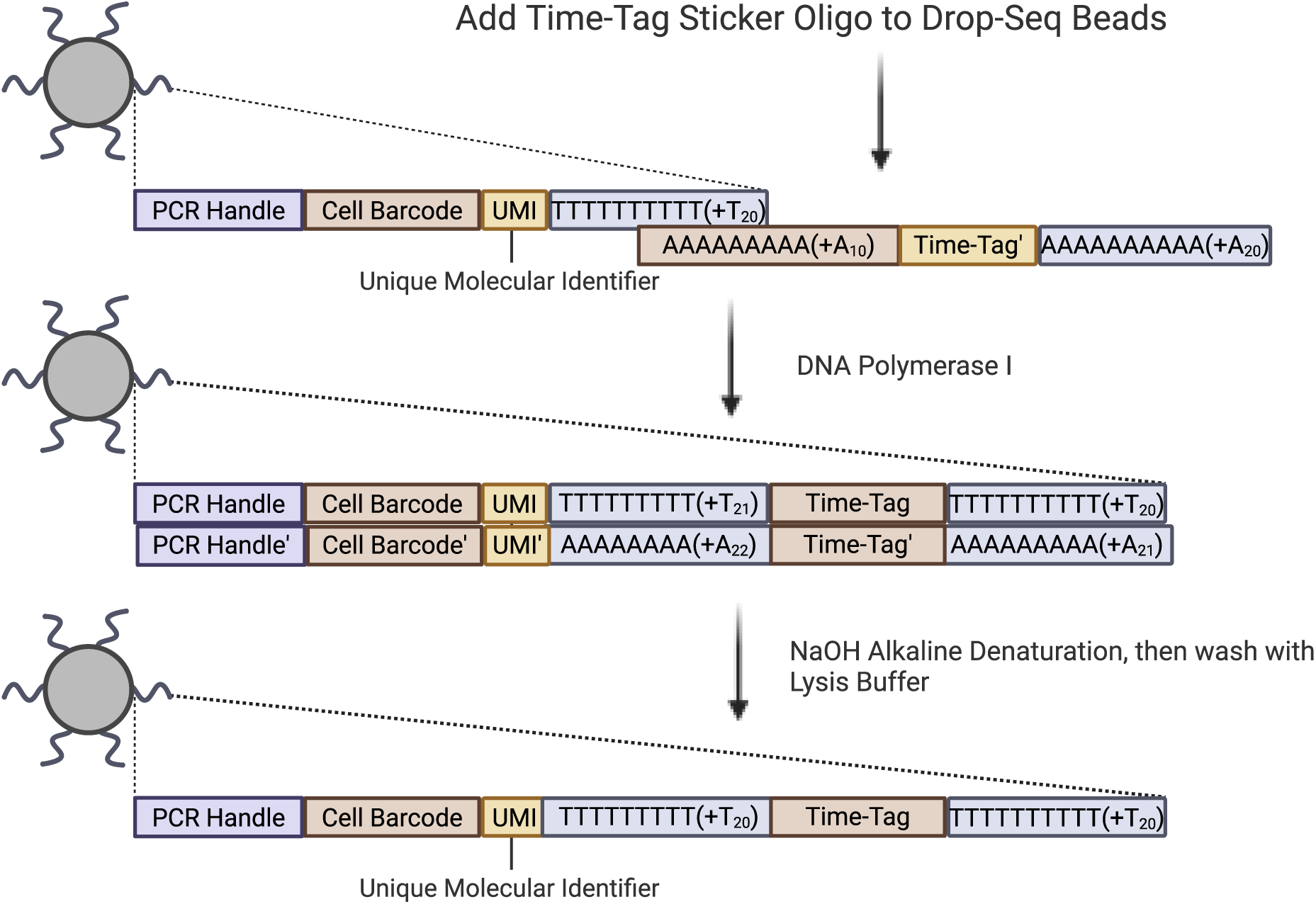
Drop-seq bead modification workflow for Creating ChronoSeq Time-Tagged beads. Sequences are added by using an oligo that binds to the existing PolyT region on Drop-seq beads. The 3’ end of the Drop-seq beads are then extended to the reverse compliment of the oligo using E. Coli DNA Polymerase I. Alkaline denaturation is used to make the double stranded DNA single-stranded. These beads are then resuspended in lysis buffer and can be used directly with our device.

**Supplementary Figure S4:**
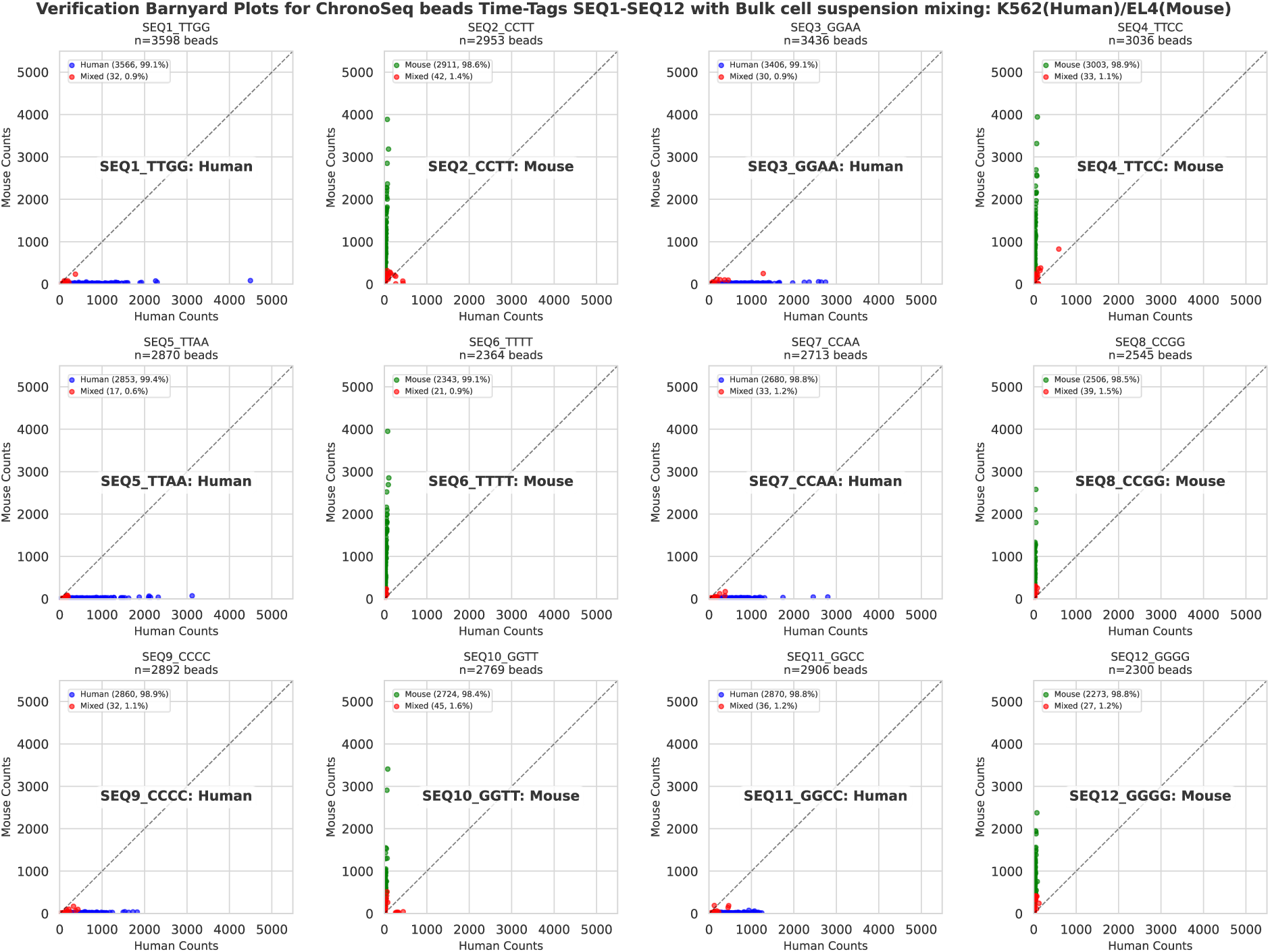
Bulk RNA-Seq Validation of all 12 Time-Tags. We checked our ability to uniquely identify each Time-Tag. We did this by mixing beads from odd numbered Time-Tags with human cells and even numbered Time-Tags with mouse cells manually in bulk. These beads were then washed and combined for reverse transcription, library preparation and sequencing. Time-Tag resolved cleanly into either human or mouse dominant with 1.6% or less mixed beads detected for each Time-Tag.10μl of Time-Tagged beads at 450 beads/μl suspended in lysis buffer were directly mixed with 8μl of mouse or human cells at 600cells/μl. Odd numbered Time-Tags were mixed with human(K562) cells while even numbered Time-Tags were mixed with mouse(EL4) cells. Each dot in the scatterplots represents a unique Cell Barcode and the (X, Y) coordinates represent the number of human and mouse transcripts captured respectively. Each plot is also labelled with the which species of the cells were mixed with the beads for the experiment.

**Supplementary Figure S5:**
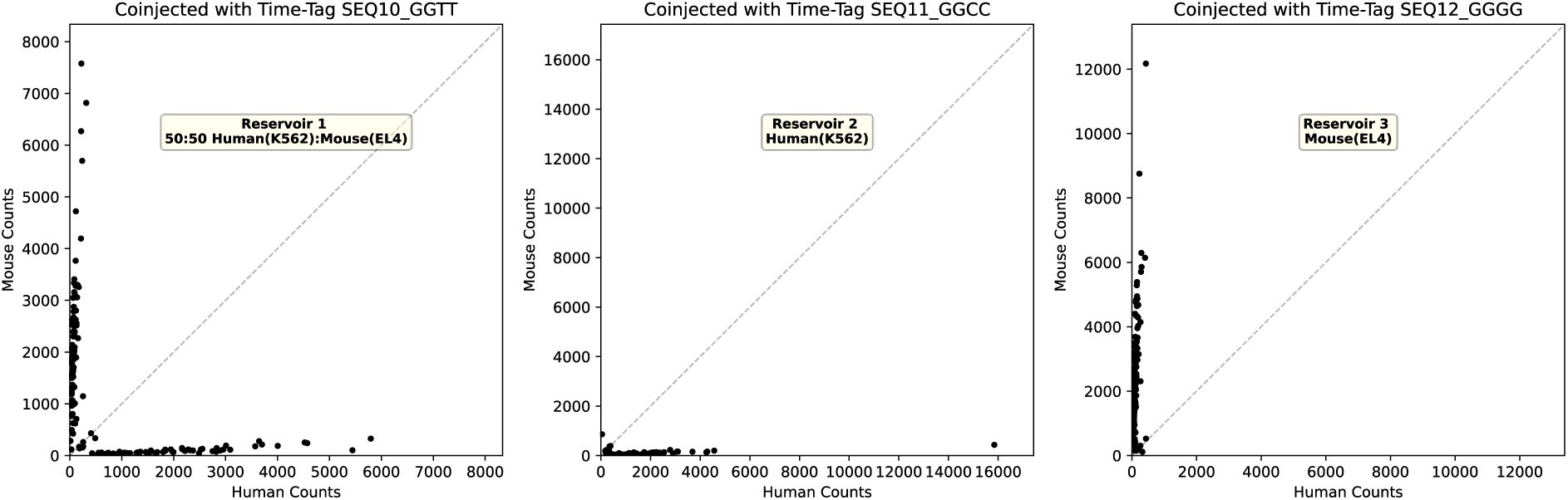
Single-Cell ChronoSeq Experiment to verify unique Time-Tagging: We checked whether the ChronoSeq device could uniquely tag each sample with a Time-Tag without mixing across samples. To do this we setup three separate reservoirs for maintaining cell suspensions inside the ChronoSeq device. Cell could be sampled from either Reservoir 1, 2 or 3 and co-injected with any of the Time-tagged beads labeled 1-12. A 50:50 mixture of human(K562) and mouse(EL4) cells was added to Reservoir 1, a human only (K562) suspension to Reservoir 2 and a mouse only(EL4) suspension to Reservoir 3. Cells were sampled with a 7-minute sampling interval including a 1-minute sampling duration using the ChronoSeq device. The first sample (left) was from a 50:50 Human(K562) and Mouse(EL4) Cell Suspension. The second sample (center) was from a Human only cell suspension. While the third sample (right) was from a Mouse only cell suspension. Each dot in the three plots represents a Unique Cell Barcode and the (X, Y) coordinates represent the number of Human and Mouse transcripts respectively. Cells were injected at a concentration of 215 cells/μl while the bead concentration was 450 beads/μl. The first, second and third cell suspensions were sequentially co-injected with Time-Tags 10 through 12. Species-mixing barnyard plots clearly showed Time-Tag 10 capturing both human and mouse cells, Time-Tag 11 capturing human cells and Time-Tag 12 capturing mouse cells.

**Supplementary Figure S6:**
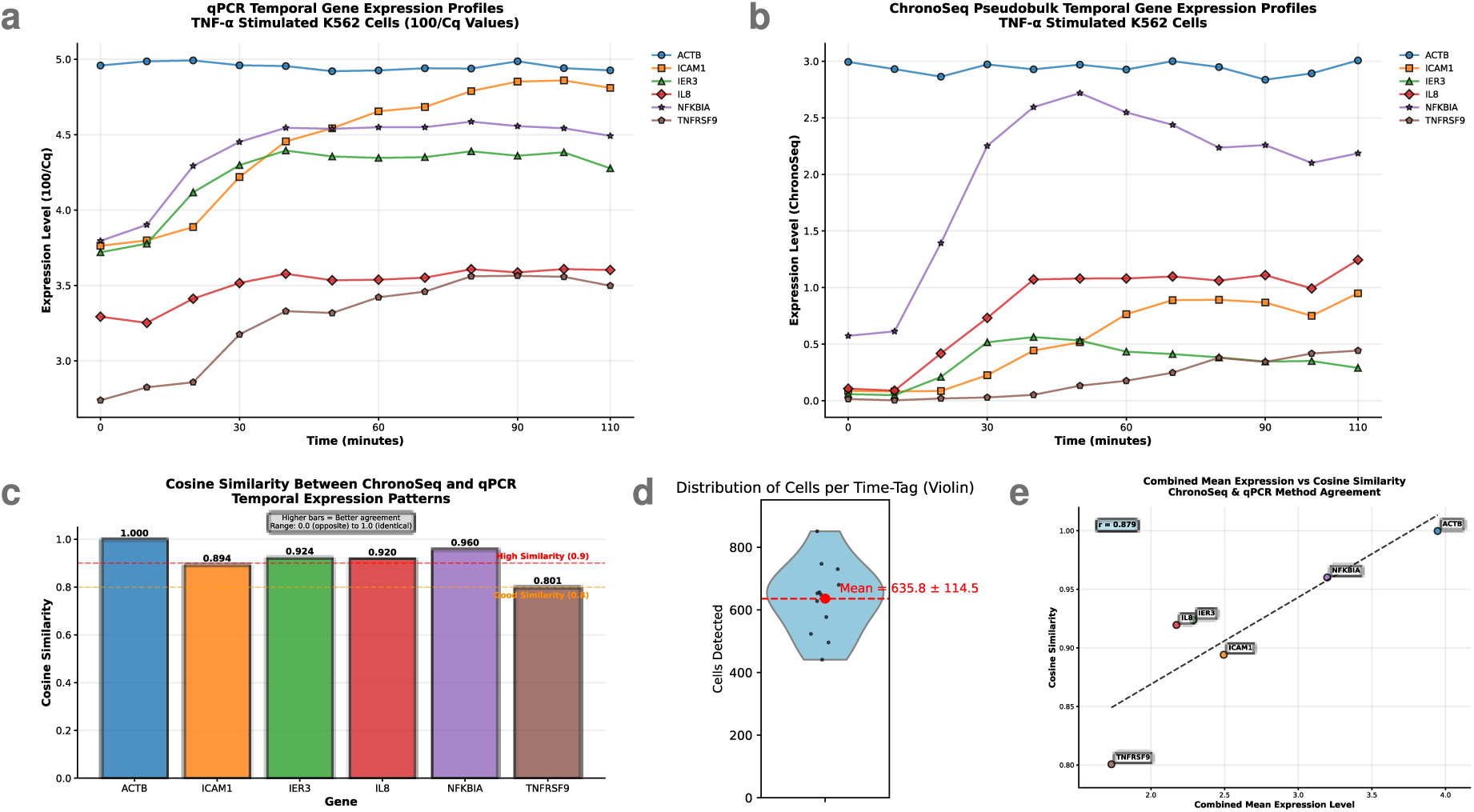
As a first demonstration of ChronoSeq, we perturbed K562 cells with 10 ng/ml TNF-α at 5 min after the first sample. We sampled every 10 minutes for 12 timepoints to capture the earliest phases of NF-κB signaling. We used a cell concentration of 430 cells/μl and a bead concentration of 450beads/μl for all Time-Tags. To validate the temporal response, we performed a qPCR measurement with the same time resolution of 10 minutes for K562 cells stimulated with TNF-α. We show 5 representative genes and one housekeeping gene (ACTB) and compare it to the pseudobulk of the same genes obtained from the single-cell ChronoSeq data. The temporal profiles (a-b) showed good to high concordance (measured by cosine similarity, c) with higher expressed genes showing higher similarity, thus less affected by noise, compared to lower expressed genes (e). We captured an average of 636 cells per Time-Tag. (d) shows a violin plot of the number of cells we captured per Time-Tag for this experiment.

**Supplementary Figure S7.**
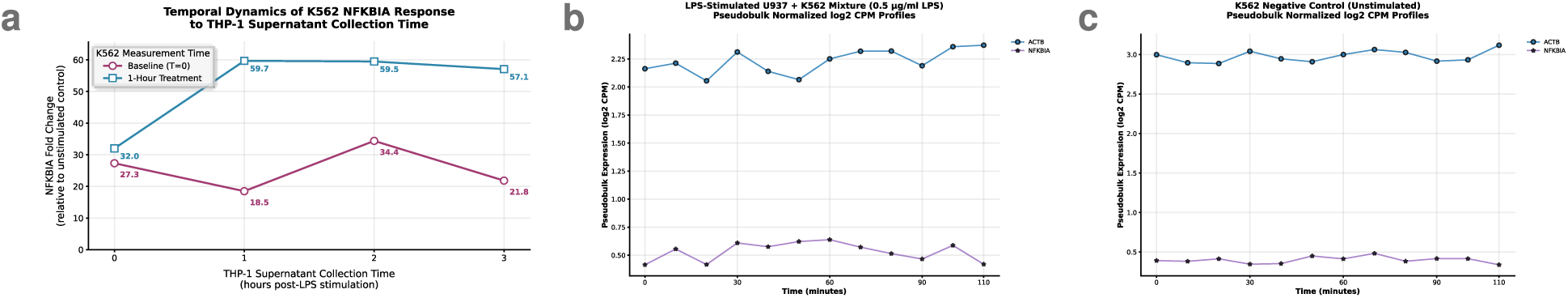
Conditioned-medium and LPS-control experiments support indirect inflammatory activation of K562 cells. We validated the lack of LPS-responsiveness for K562 cells with a qPCR experiment where supernatants from THP-1 cells stimulated with LPS were added to K562 cells at different times after LPS stimulation. NFKBIA was measured as a marker of the inflammatory response in K562 cells and ACTB was measured as a control. K562 supernatants show strong activation for supernatants taken after 1 hour or longer from THP-1 cells: (a) NFKBIA fold changes were calculated using the 2^(-ΔΔCq)^ method^73^ with ACTB as the housekeeping gene and unstimulated K562 cells as the reference control. Temporal dynamics were visualized by plotting fold change values against THP-1 supernatant collection time (0, 1, 2, 3 hours post-LPS stimulation). Two K562 measurement timepoints were compared: baseline (T=0, immediately after supernatant addition) and 1-hour post-treatment (T=1h). Data points represent mean values from n=2 technical replicates. Plots clearly show K562 response to stimulation with THP-1 supernatants collected at 1 hour with double the fold change. To further show that K562 cells cannot respond to LPS either alone or when mixed with cells unresponsive to LPS; we LPS stimulated a 50:50 mixture of K562 and undifferentiated U937 cells which are known to show limited or no response to LPS^74^. Unlike the pseudobulks of NFKBIA and ACTB in K562 cells stimulated with TNF-α (**Fig. S6(b)**), the U937:K562 mixture shows no response: (b) Pseudobulks for scRNA-seq generated for ACTB and NFKBIA genes from a 50:50 U937 and K562 cells mixture stimulated with 0.5ug/ml LPS inside the ChronoSeq device. Samples were taken every 10 minutes for 12 timepoints, with LPS addition 5 minutes after first sample. This lack of response is similar to NFKBIA and ACTB pseudobulks from the K562 negative control experiment: (c) Pseudobulks for scRNA-seq generated for ACTB and NFKBIA genes from the K562 negative control experiment.

**Supplementary Figure S8:**
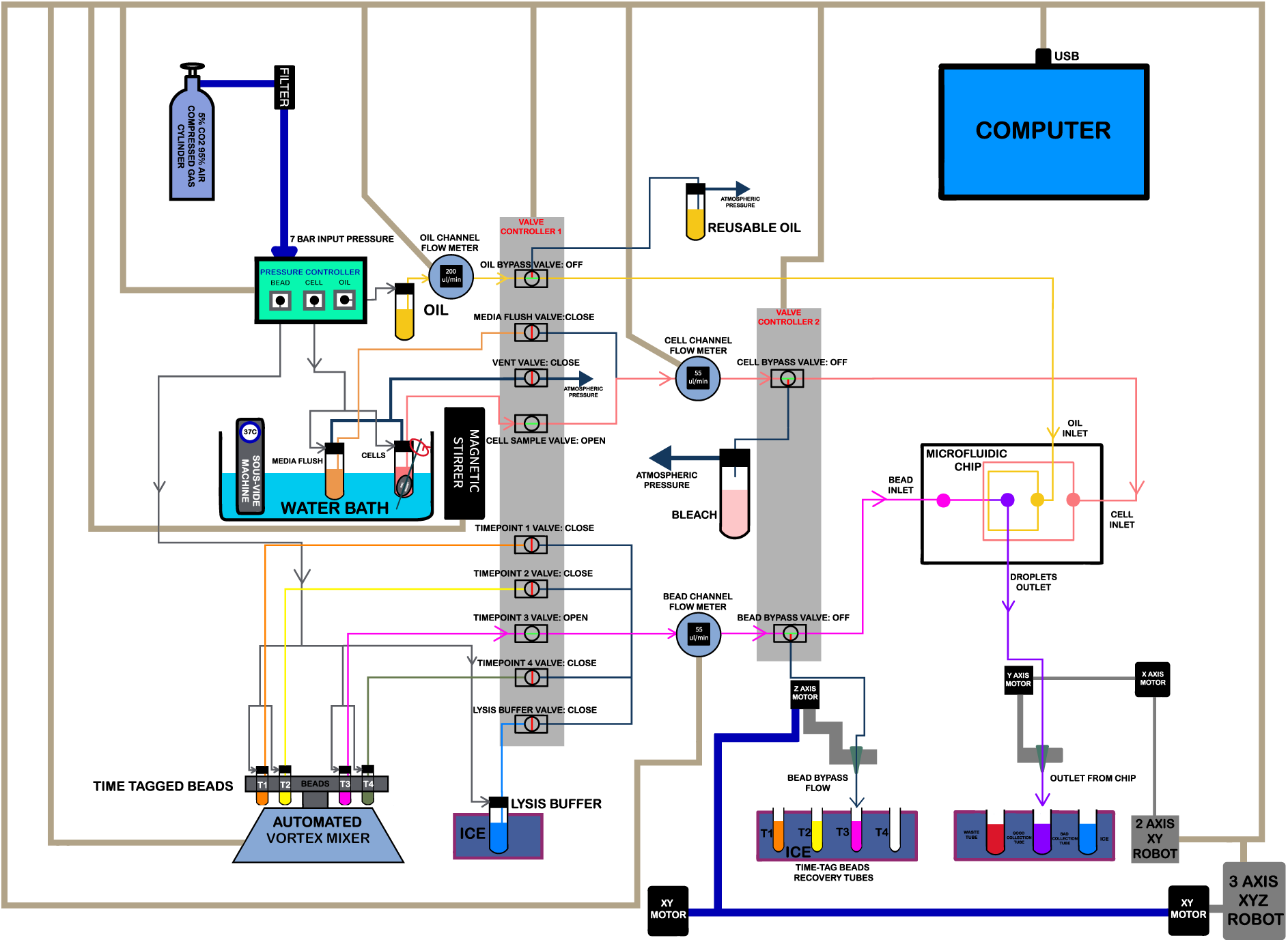
ChronoSeq device full schematic.

**Supplementary Figure S9:**
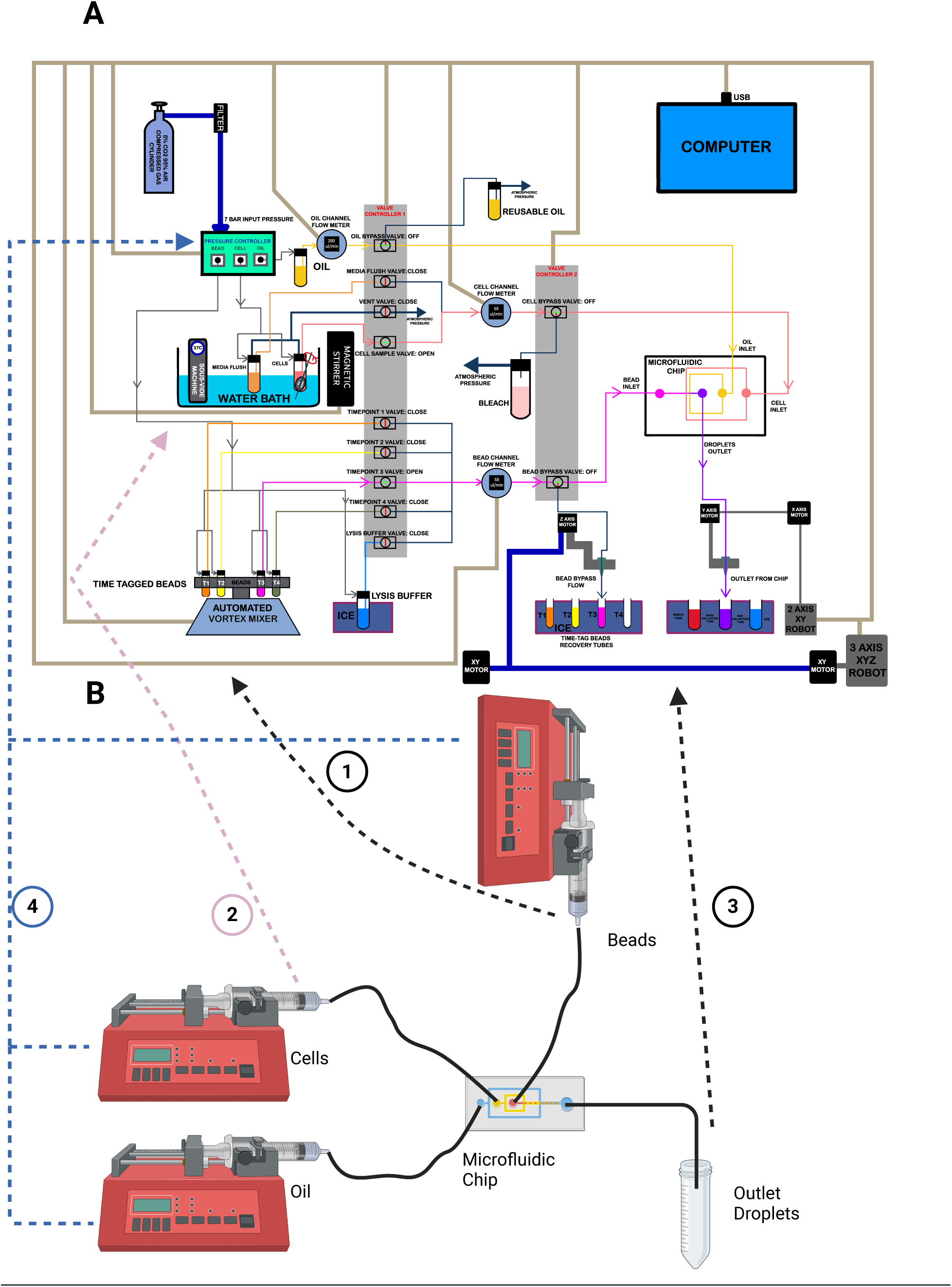
Comparison of ChronoSeq and Drop-seq devices. Our device design addresses several challenges not present in the original Drop-seq^52^ system. First, extended cell viability. We maintain cell suspension viability for experiments lasting 2 hours or longer, compared to typical 30-minute Drop-seq runs. Second, cross-contamination prevention. We implemented a washing system to remove unused cells or beads between samples. Third, sample isolation. We designed a system to prevent mixing wash fluid with droplets. To address these challenges, our device operates in two main phases during each injection cycle. 1. Flushing phase that clears fluid lines of residual beads or cells from previous injections. 2. Injection phase that precisely introduces cells and Time-Tagged beads into the microfluidic chip. This integrated system enables controlled, sequential introduction of cells and Time-Tagged beads, capturing gene expression dynamics across multiple time points in a single experiment. Our design prioritizes cell viability maintenance, precise fluid control, and efficient bead usage, all critical for generating high-quality, time-resolved single-cell sequencing data. The ChronoSeq device is controlled via serial connections to a computer, which coordinates the actions of its various components, while the Drop-seq device works by setting flow rates manually on syringe pumps. Some key features distinguish the ChronoSeq device:

1. **Bead handling:** ChronoSeq incorporates an automated vortex mixer to keep the beads evenly suspended before injection. Drop-seq uses a magnetic stirrer for bead suspension, which leads to high dead volumes and complicates automation for multiple Time-Tags. ChronoSeq allows for the easy addition of more Time-Tags in the future if necessary. Additionally, loading and unloading the beads is simplified; we can directly screw a new 50 mL tube containing beads into the reservoir. The Time-Tagged bead reservoirs are connected to timepoint valves, which facilitate the selection of the appropriate Time-Tagged beads for injection into the microfluidic chip.
2. **Cell culture system:** In the Drop-seq device, cells are directly loaded into a syringe for injection, typically running for 15 to 30 minutes. However, our goal is to maintain cell viability for at least 2 hours. Leaving cells at room temperature without growth media and in the absence of 5% CO_2_ can lead to excessive cell death and unwanted perturbations. To address this, we have integrated a cell culture system directly into the ChronoSeq device, featuring a water bath to maintain cells at 37°C, along with a 5% CO_2_ gas cylinder that serves as a pressure source and aids in cell culture. Furthermore, we can add additional tubes to culture multiple cell suspensions separately and use valves to select which cell suspension is injected into the microfluidic chip.
3. **Droplet collection:** In the Drop-seq device, droplets are collected directly in a 50 mL tube kept on ice. However, in our system, it is crucial to prevent the beads and cells from previous injections from mixing with those from new injections. To achieve this, we developed a washing system to clear the fluid lines of residual beads and cells. We also need to ensure that washing liquids and unused cell suspensions do not mix with the collected droplets. To solve this issue, we designed a robotic system that facilitates droplet collection and bead recovery. This robot automatically moves the outlet tubing to the correct tube, preventing the mixing of droplets with washing fluids. Additionally, it can recover unused beads from the fluid lines, minimizing waste and reducing experimental costs. Like the Drop-seq device, the droplets in our system are also kept on ice using ice boxes.
4. **Flow control:** In the Drop-seq device, precise speed control is essential for droplet formation. Speeds that are too high can inhibit droplet formation, while low speeds can negatively impact throughput. To achieve this precise control, Drop-seq employs syringe pumps, which allow direct manipulation of the syringe’s squeezing rate. In our case, we require high flow rates for flushing and low flow rates for droplet formation. Low flow rates like Drop-seq flow rates, are insufficient for quickly washing unused beads and cells from the device. Therefore, we implemented a compressed air pressure controller and flow sensors for each of the bead, cell, and oil channels. The compressed air controller enables rapid changes in flow rate, while the flow sensors provide the precise control needed for high-quality droplet formation during low flow rate operation.

**Supplementary Figure S10.**
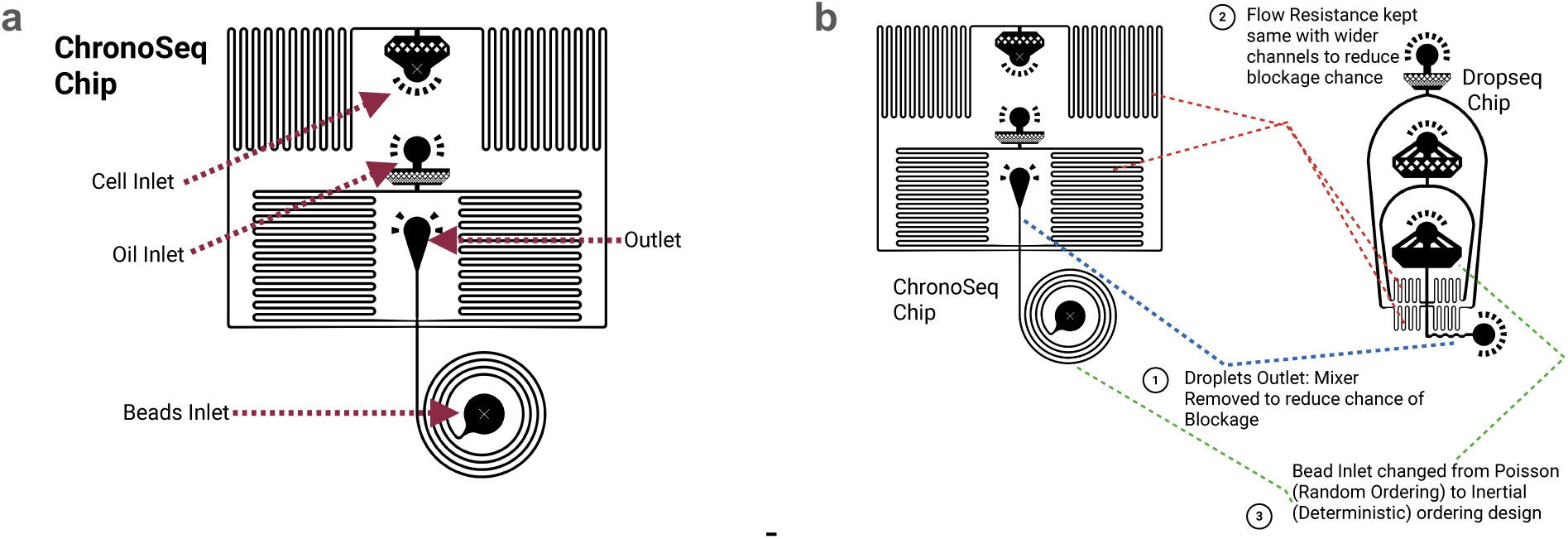
ChronoSeq microfluidic chip. The primary function of the microfluidic chip is to co-encapsulate cells with Time-Tagged beads using a flow-focusing junction^75^ at the center of the chip. While we initially considered using the Drop-seq device for this purpose, several limitations made it unsuitable for our application: 1. Ordering mechanism. Drop-seq relies on random (Poisson) ordering for co-encapsulation of cells and beads. This random ordering makes it difficult to increase the number of cells collected per timepoint without also increasing droplets containing multiple cells or beads. The ChronoSeq microfluidic chip uses an optimized version of an existing inertial ordering design^76^ to increase cell capture through deterministic ordering of beads into droplets. This deterministic ordering allows for capturing more cells by increasing the number of droplets containing exactly one bead and one cell. 2. Channel design. Both the Drop-seq chip and existing inertial ordering^76^ designs use narrow flow resistance channels, which are prone to blockage during the flushing phase. To reduce blockage risk, we widened the channels in our chip design. To maintain the same flow resistance with wider channels, we increased the length of the channels to compensate. **(a) ChronoSeq chip design** with labeled inlet and outlet holes. **(b) Comparison of the chip design**. The ChronoSeq chip has three key modifications compared to the Drop-seq chip. 1: The Droplet Mixer near the Drop-seq outlet hole has been removed to reduce the likelihood of blockage. 2: The width of the flow-resistors has been increased along with the length of the channels to keep the flow resistance identical between the two designs. 3. The straight (Poisson) ordering inlet has been replaced with an inertial (deterministic) ordering spiral design. These modifications aim to improve the chip’s performance in terms of reduced clogging, maintained flow dynamics, and enhanced cell/bead ordering for more efficient single-cell encapsulation. The inertial ordering design, in particular, allows for a more deterministic approach to cell and bead pairing, potentially increasing the capture efficiency of single cells with single beads.

**Supplementary Figure S11.**
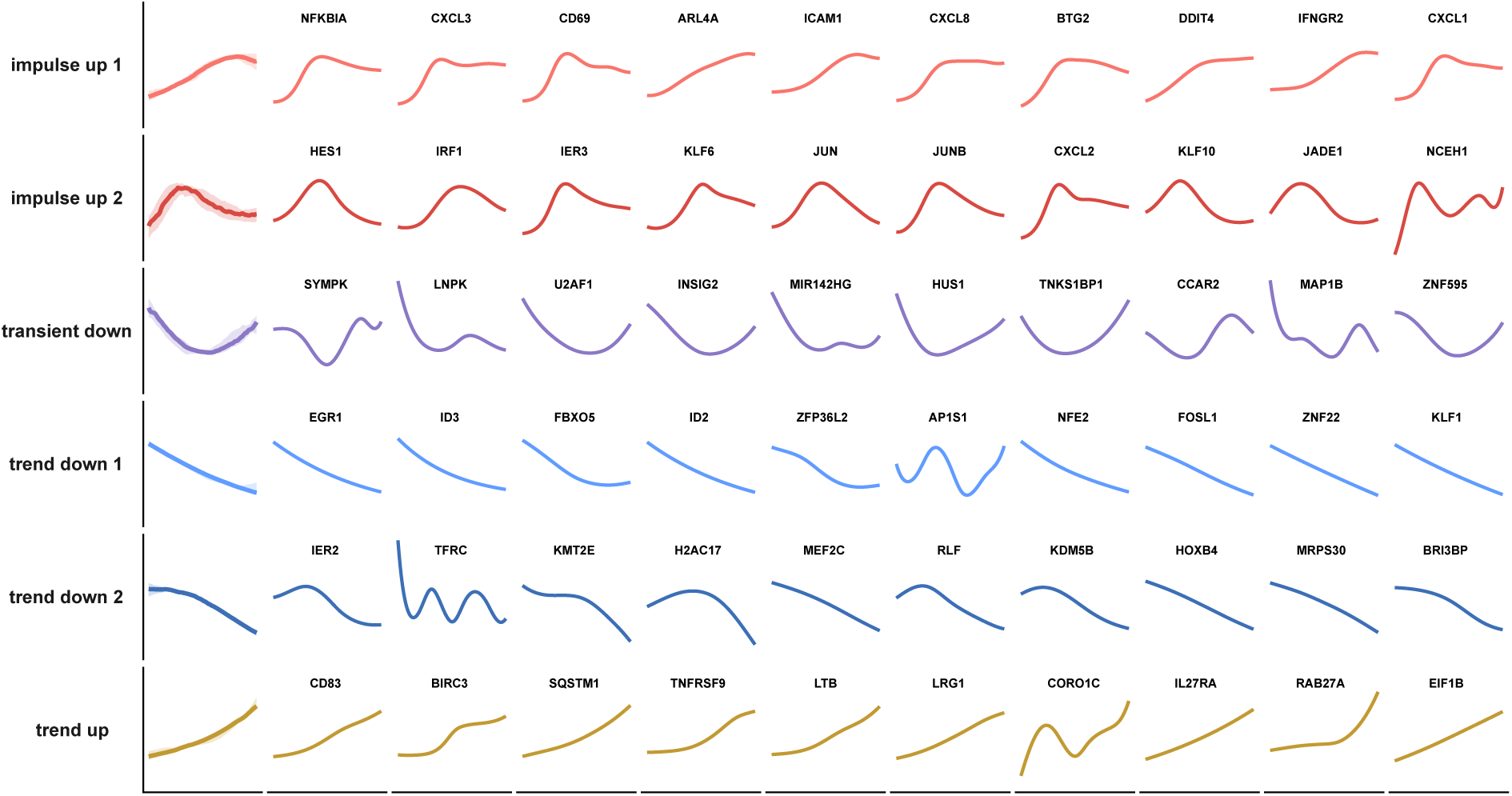
K562+TNF-α dynamic response modes with (*left*) median curves and (*right*) representative genes in each cluster.

**Supplementary Figure S12.**
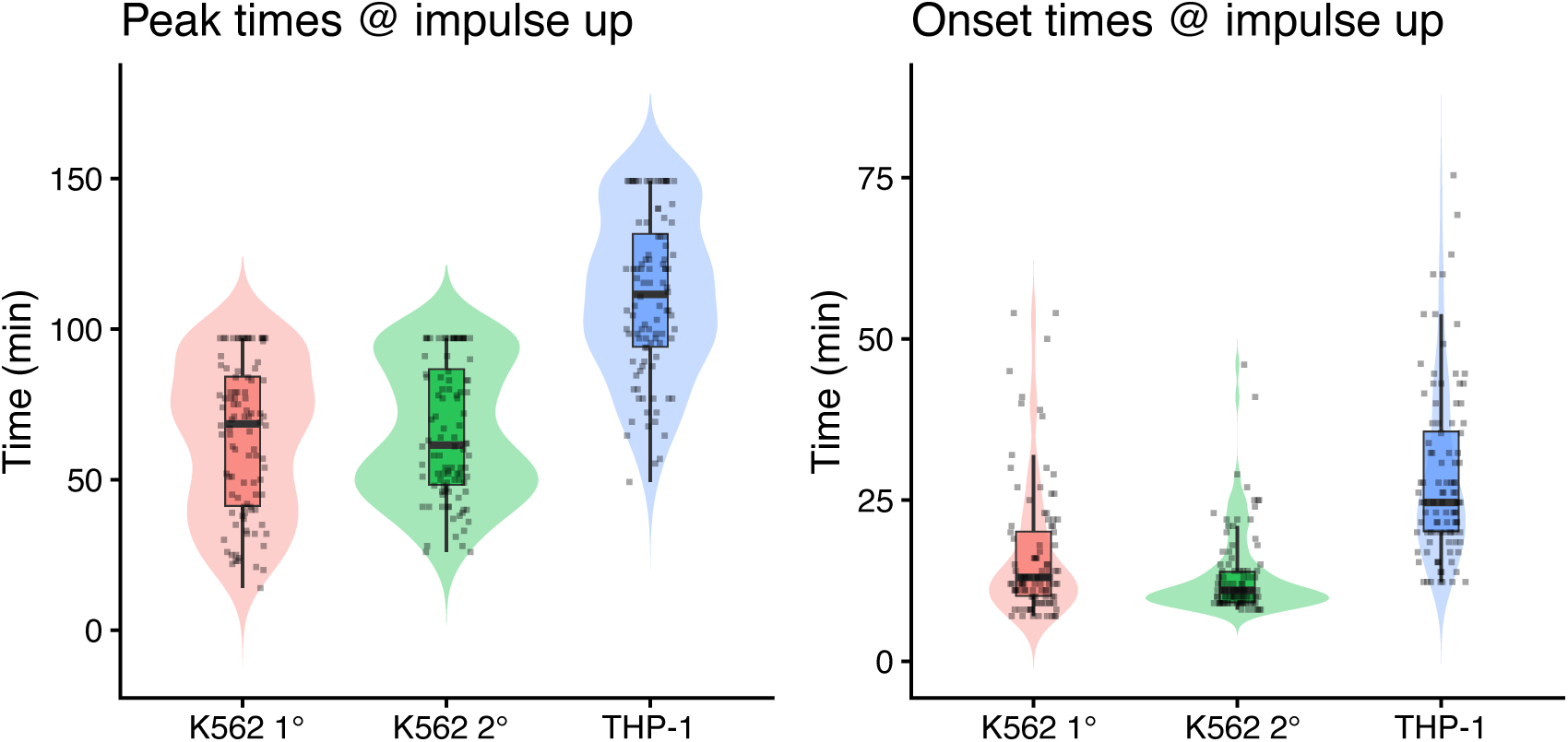
Onset and peak times of impulse-up genes across inflammatory contexts. Distributions of estimated peak (*left*) and onset (*right*) times for both impulse-up subtypes, calculated after aligning transcriptional responses across contexts using cell population-specific offsets (see **Methods**). K562 1° = K562 + TNF-α. THP-1 = THP-1 + LPS. K562 2° = K562 response in co-culture with LPS-stimulated THP-1 cells.

**Supplementary Figure S13.**
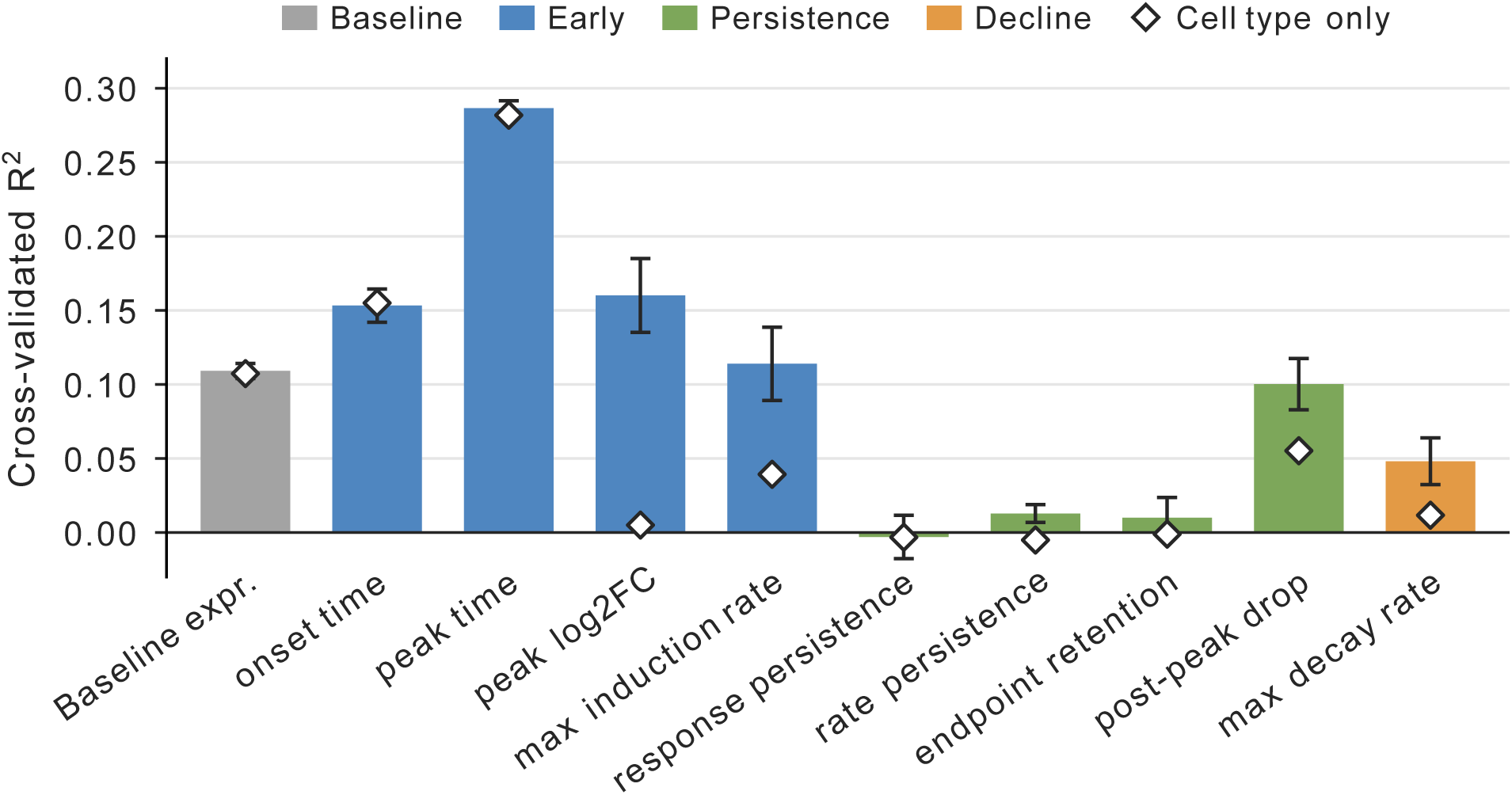
Cross-validated predictive performance of regulatory features across continuous gene expression kinetics. Bars show the mean cross-validated R² of full elastic net models incorporating promoter architecture, enhancer features, RNA stability, gene length, and cell type, while diamonds indicate performance using cell type alone. Models were evaluated using repeated nested cross-validation grouped by gene. Colors distinguish descriptors of baseline expression, early response kinetics, persistence, and decline. Error bars indicate SD across cross-validation repeats.

**Supplementary Figure S14.**
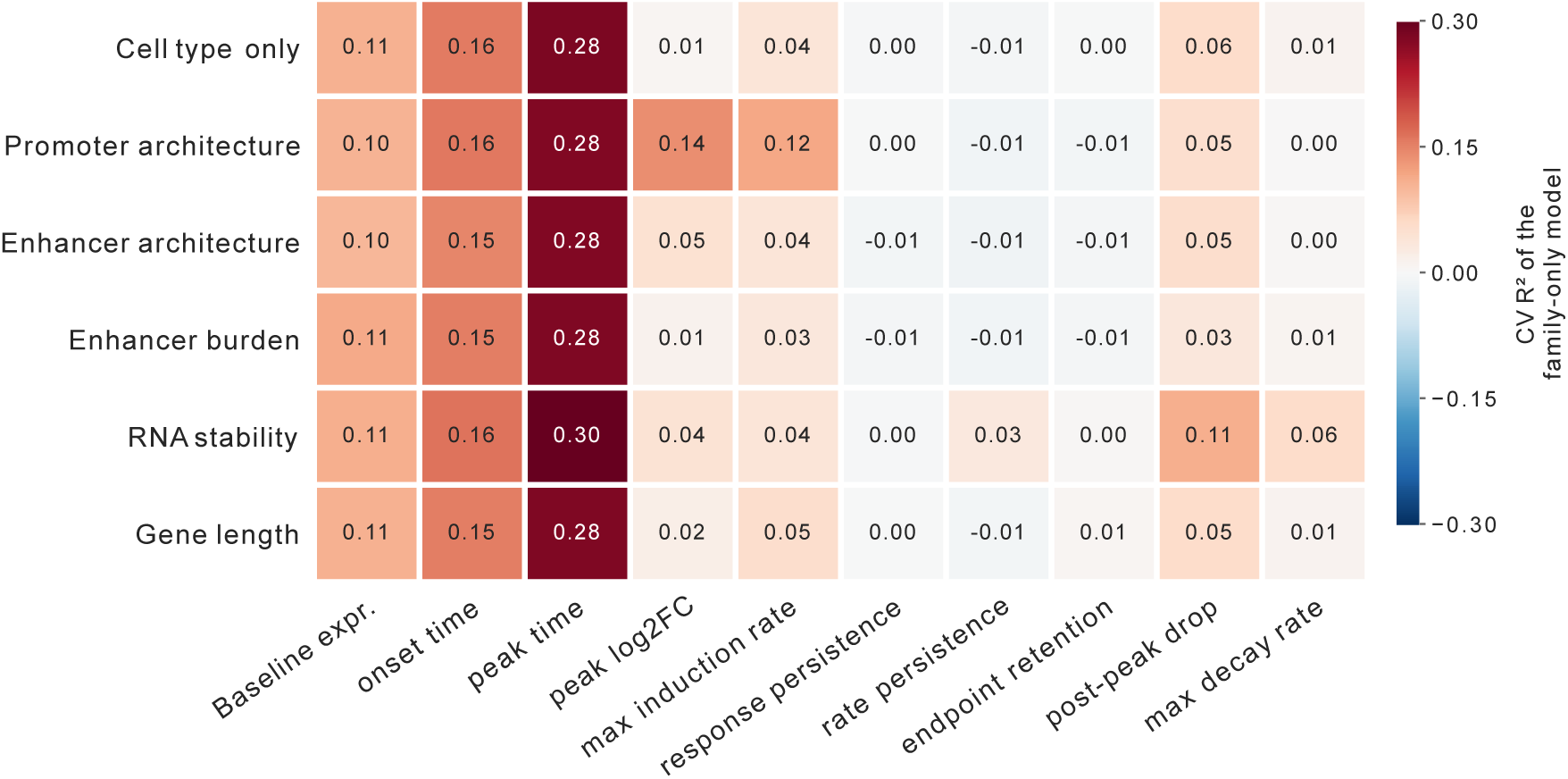
Cross-validated predictive performance of individual feature families across continuous gene expression kinetics. Heatmap shows the mean cross-validated R² of elastic net models incorporating cell type alone or cell type together with individual feature families, including promoter architecture, enhancer architecture, enhancer burden, RNA stability, and gene length. Models were evaluated using repeated nested cross-validation grouped by gene. Colors indicate predictive performance, with values shown in each cell.

**Supplementary Figure S15.**
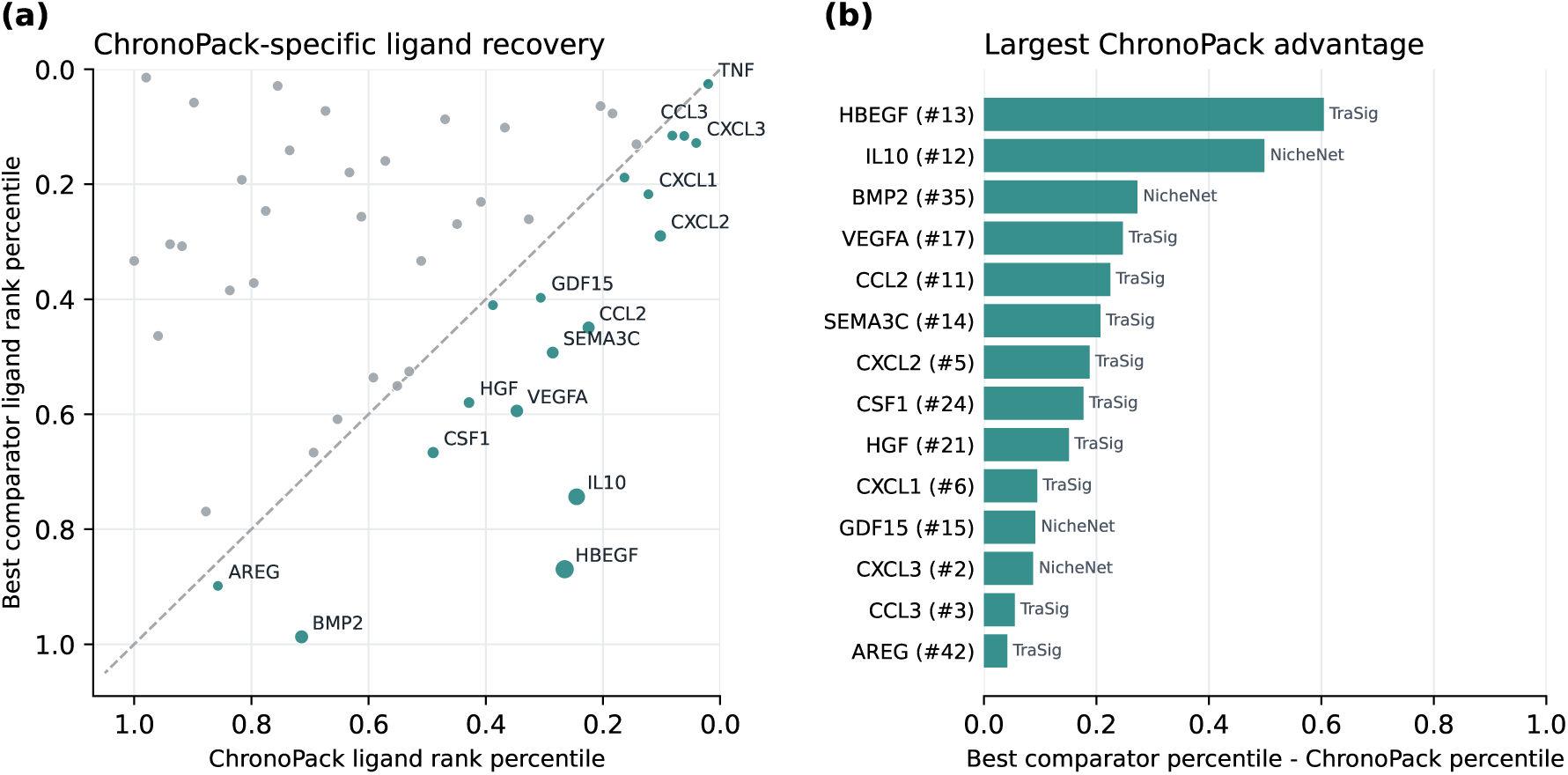
ChronoPack prioritizes candidate signaling ligands ranked lower by existing methods. Ligand rankings for THP-1–K562 cell–cell communication were compared between ChronoPack and four methods: LIANA, CellChat, NicheNet, and TraSig. Comparator ligand ranks were determined by their highest-scoring ligand–receptor pair, with ties assigned mean ranks. Rankings were converted to percentiles within each method’s candidate ligand set, with lower values indicating higher priority. **(a)** Comparison of ChronoPack rank percentiles with the best percentile across comparators. Teal points represent ligands ranked higher by ChronoPack than by all comparators, with point size reflecting the ranking difference. **(b)** The 14 ligands showing the largest ChronoPack advantage, defined as the best comparator percentile minus the ChronoPack percentile. Parentheses indicate ChronoPack ranks, and bar labels identify the highest-ranking comparator.

**Supplementary Figure S16.**
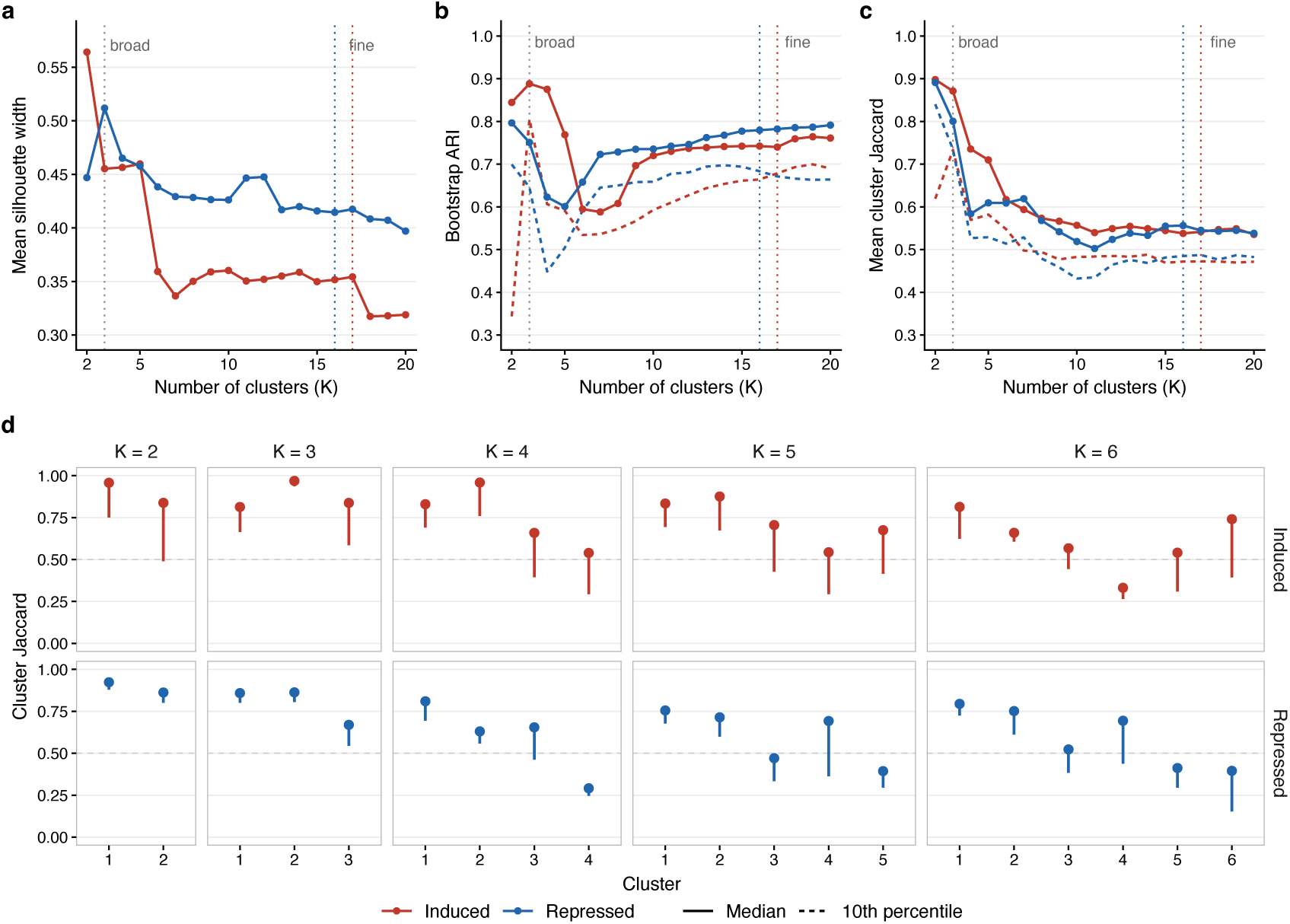
Resolution and stability of gene trajectory clustering. **(a)** Mean silhouette width of induced and repressed gene partitions across K = 2–20 clusters. **(b, c)** Clustering stability assessed using 200 random subsamples, each retaining 80% of gene trajectories. Subsample-derived partitions were compared with the corresponding full-data partitions restricted to the same trajectories, using the adjusted Rand index (**b**) and mean best-match Jaccard index across full-data clusters **(c)**. Solid and dashed lines indicate the median and 10th percentile across subsamples, respectively. Vertical dotted lines mark the broad and fine clustering resolutions. **(d)** Cluster-level stability at K = 2–6 for induced and repressed trajectories, quantified by the best-match Jaccard index for each full-data cluster. Points indicate medians across subsamples, with vertical lines extending to the 10th percentile.

**Supplementary Table S1:** Cq Values for HPRT1 for K562 cells stimulated THP-1 supernatant and controls: NaN means no Cq value obtained due to lack of amplification for that sample. Results are consistent across both replicates and show addition of THP-1 supernatant leads to RNA degradation.

| Gene | Stimulant for K562 cells | K562 cells stimulation duration | Replicate | Cq |
|---|---|---|---|---|
| HPRT1 | LPS stimulated THP-1 cells supernatant taken at 2 hours | 1 hour | 1 | NaN |
| HPRT1 | LPS stimulated THP-1 cells supernatant taken at 2 hours | 1 hour | 2 | NaN |
| HPRT1 | LPS stimulated THP-1 cells supernatant taken immediately-0 hours | 1 hour | 1 | NaN |
| HPRT1 | LPS stimulated THP-1 cells supernatant taken immediately-0 hours | 1 hour | 2 | NaN |
| HPRT1 | Cell culture media DMEM- No stimulation | 1 hour | 1 | 32.48 |
| HPRT1 | Cell culture media DMEM-No stimulation | 1 hour | 2 | 32.29 |
| HPRT1 | 10ng/ml TNF-α | 1 hour | 1 | 31.88 |
| HPRT1 | 10ng/ml TNF-α | 1 hour | 2 | 31.68 |
| HPRT1 | Blank Control no K562 cells only water | N/A | 1 | NaN |
| HPRT1 | Blank Control no K562 cells only water | N/A | 2 | NaN |

